# Fc receptor dependent and independent mechanisms of antibody-mediated enhancement of immune responses

**DOI:** 10.64898/2026.03.27.714269

**Authors:** Melissa Cipolla, Andrew J. MacLean, Brianna Hernandez, Gabriela S. Silva Santos, Leonidas Stamatatos, Anna Gazumyan, Harald Hartweger, Julia Merkenschlager, Stylianos Bournazos, Jeffrey V. Ravetch, Michel C. Nussenzweig

## Abstract

Immune memory responses are rapid and qualitatively distinct from primary responses. They typically develop in the presence of antigen-experienced memory T and B cells and pre-existing antibodies. Although the contribution of T and B cells to recall responses is well defined, the contribution of antibody “memory” and the mechanisms by which pre-existing antibodies modulate the development of germinal center and plasma cell responses is not precisely understood. Here we report on mechanisms that mediate antibody enhancement of germinal center (GC) and plasmablast (PB) compartments, and the parallel process by which they change the affinity threshold for B cell recruitment into immune responses. The data indicate that antibody-mediated enhancement of GC and PB responses is Fc gamma receptor (FcγR) dependent and largely complement receptor 1 and 2 (CR1/2) independent. In contrast, the reduction in the affinity threshold for GC entry is independent of both FcγRs and CR1/2.

**Summary:** Cipolla et al. show that antibody can modulate immune responses via both Fc gamma receptor dependent and independent mechanisms. These mechanisms influence both the magnitude and composition of the germinal center response.

## Introduction

Recall immune responses form the basis of vaccine-mediated protection from pathogens. These responses involve both cellular and humoral components, T and B lymphocytes and antibodies, respectively. Exposure to antigens or pathogens induces CD4 and CD8 T cell proliferative expansion accompanied by changes in gene expression that poise these cells to generate a more rapid and effective response upon subsequent re-exposure (Weng et al., 2012; Künzli and Masopust, 2023). Similarly, B cells recruited to GCs during primary immune responses produce memory compartments containing B lymphocytes which upon re-exposure can also respond rapidly (Weisel and Shlomchik, 2015; Inoue and Kurosaki, 2024; Ripperger and Bhattacharya, 2021). Finally, antibodies produced by plasma cells during the primary immune response constitute additional memory that can block or neutralize pathogens (Bournazos and Ravetch, 2017).

In experiments performed during the early 20^th^ century, Smith observed that pre-existing antibodies to diphtheria toxin can suppress endogenous immune responses (Smith, 1909). Subsequent work focused on dissecting the intricacies of antibody-mediated suppression, while also elucidating a role for antibodies in enhancing immunity (Finkelstein and Uhr, 1964; Henry and Jerne, 1968; Walker and Siskind, 1968; Murgita and Vas, 1972; Pincus and Nussenzweig, 1969; Brüggemann and Rajewsky, 1982; Enriquez-Rincon and Klaus, 1984; Wiersma et al., 1989; Wiersma, 1992). Experiments to dissect the role of an antibody’s Fc domain in these phenomena and its interactions with FcγRs and the complement cascade produced conflicting results that were further confounded by the use of supraphysiological doses of highly heterogenous antigens like sheep red blood cells with unknown epitope specificity or model hapten antigens with single dominant epitopes (Tao and Uhr, 1966; Sinclair et al., 1968; Sinclair, 1969; Heyman, 1989; Wiersma et al., 1990; Wernersson et al., 1999; Getahun et al., 2004; Ståhl and Heyman, 2001).

More recent work on SARS-CoV-2 vaccinated human volunteers and BCR (B-cell receptor) knock-in mice infused with specific monoclonal antibodies showed that pre-existing antibodies can block immunodominant epitopes and redirect GC B cell responses to subdominant epitopes (McNamara et al., 2020; Schaefer-Babajew et al., 2023; Hägglöf et al., 2023; Dvorscek et al., 2024; Tas et al., 2022). In particular, enhancement was associated with low affinity antibodies and masking with high affinity interactions (Tas et al., 2022; Dvorscek et al., 2024). However, the precise mechanisms by which antibodies mediate enhancement, suppression and diversification of polyclonal immune responses to complex antigens in wild-type (WT) mice remains to be determined. Here we show that antibodies enhance polyclonal immune responses by at least two distinct mechanisms. The first is FcγR dependent and increases the magnitude of GC and PB responses. The second is FcγR independent and influences the affinity threshold and thereby the diversity of B cells seeding the immune response.

## Results

### Antibody enhancement of immune responses

To examine how antibodies modulate the development of immune responses in the absence of the potentially confounding effects of cellular memory, we infused WT mice with an anti-HIV-1 antibody 3BNC117 IgG2a and immunized them the following day with TM4-Core, an HIV-1 envelope immunogen (Dosenovic et al., 2018; McGuire et al., 2014; Scheid et al., 2011) (Fig. 1 A). The variable region of 3BNC117 IgG2a binds to the CD4 binding site of the HIV-1 envelope and its murine Fc region binds to both activating and inhibitory FcγRs (Bournazos et al., 2014). Seven days after immunization, 3BNC117 IgG2a infused mice showed significantly larger GC and PB responses in the draining lymph nodes (LNs) compared to controls (Fig. 1, B-G; and Fig. S1 A). However, the fraction of GC B cells that bound to antigen with sufficient affinity to be detected by flow cytometry using HIV-1 envelope tetramers was significantly reduced in antibody infused mice (Fig. 1, H-I; and Fig. S1 A). Despite the relative reduction in frequency, the absolute number of antigen-binding cells was similar in the two experimental groups, indicating that pre-existing antibody can recruit additional B cells to the response S1 A). Thus, antibody can both enhance and diversify the repertoire of B cells participating in the GC reaction.

**Figure 1:**
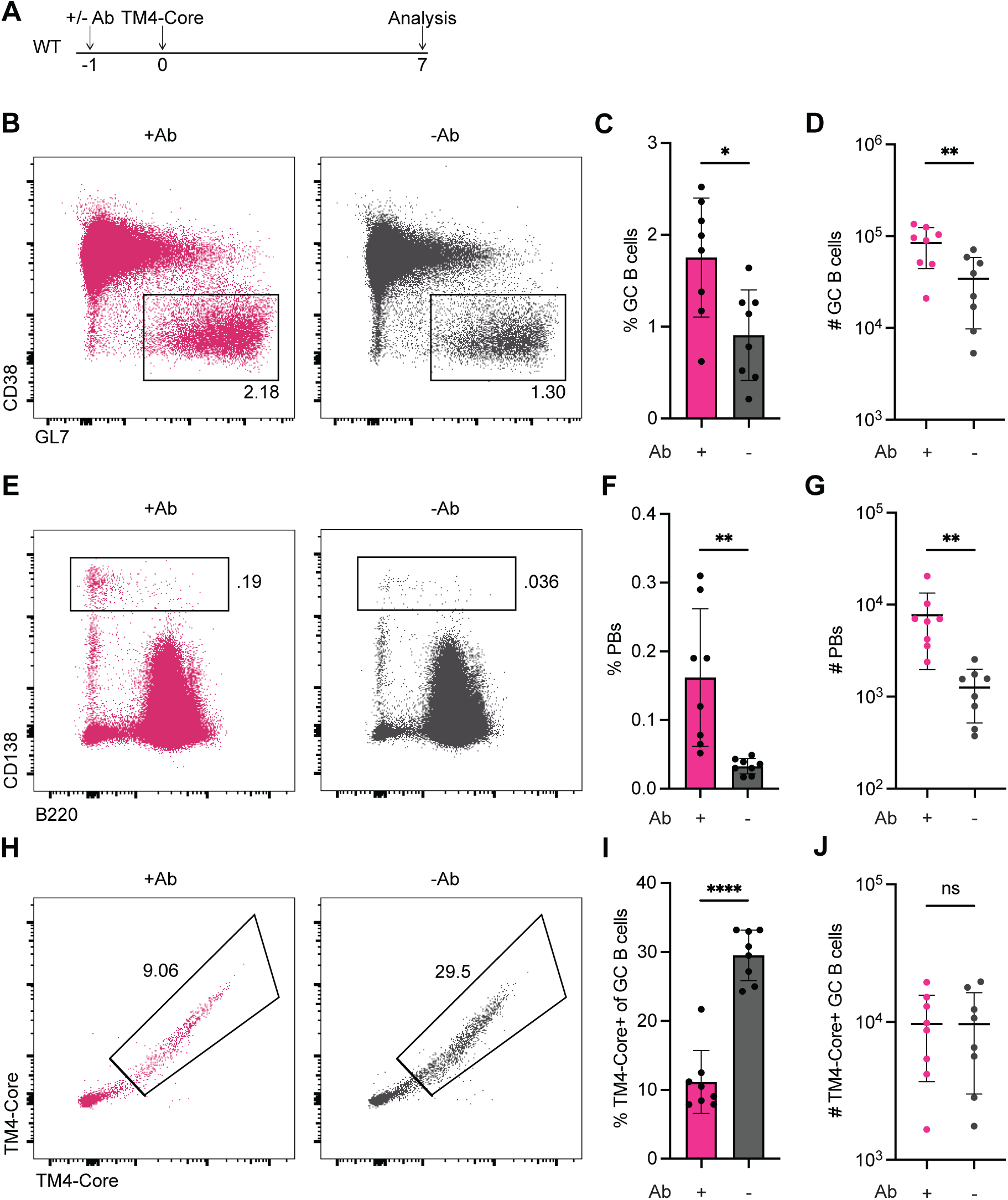
Antibody modulates immune responses in the absence of cellular memory. **(A)** Experimental schematic for **B-J**. (**B)** Representative flow cytometry plots for GC B cells in mice pre-infused with 3BNC117 IgG2a (left, +Ab, pink, for **B-J**) or no antibody (control, right, -Ab, grey, for **B-J**). GC B cells are further gated on CD95+. **(C)** Plot shows the percentage of GC B cells of B220+ cells in mice pre-infused with 3BNC117 IgG2a or control. **(D)** Plot shows the number of GC B cells in mice pre-infused with 3BNC117 IgG2a or control. **(E)** Representative flow cytometry plots for PBs in mice pre- infused with 3BNC117 IgG2a or control. **(F)** Plot shows the percentage of PBs in mice pre-infused with 3BNC117 IgG2a or control. **(G)** Plot shows the number of PBs in mice pre-infused with 3BNC117 IgG2a or control. **(H)** Representative flow cytometry plots for TM4-Core+ GC B cells in mice pre-infused with 3BNC117 IgG2a or control. **(I)** Plot shows the percentage of TM4-Core+ cells of GC B cells in mice pre-infused with 3BNC117 IgG2a or control. **(J)** Plot shows the number of TM4-Core+ GC B cells in mice pre-infused with 3BNC117 IgG2a or control. Data is displayed as mean ± SD. Each point represents a single mouse from two independent experiments. Statistical significance for **C, D, F, G, I** and **J** was determined using two-tailed unpaired t tests where ns denotes non-significant, * denotes p ≤ 0.05, ** denotes p ≤ 0.01 and **** denotes p ≤ 0.0001.

To rule out the possibility that these phenomena were HIV-1 TM4-Core specific, we repeated the experiments using SARS-CoV-2 Spike and C144 IgG2a, an antibody which binds the SARS-CoV-2 receptor binding domain (RBD) (Fig. S2 A). Pre-infusion with C144 IgG2a produced larger GC and PB compartments compared to controls S2 B-E). Moreover, the GCs in antibody infused mice contained a reduced fraction of B cells that bound to the RBD; however, the absolute numbers of antigen-binding cells were unchanged due to overall larger GC reactions (Fig. S2 F and G). The data indicates that the antibody-mediated enhancement and diversification of humoral immune responses is not limited to HIV-1 TM4-Core.

### T cells in antibody-mediated enhancement

To determine whether the increased magnitude of the GC and PB compartments is dependent on T cell help, we treated mice with anti-CD40L or an isotype control on days 3 and 6 after immunization (Fig. 2 A). Compared to mice receiving an isotype control, 3BNC117 IgG2a infused mice receiving anti-CD40L had significantly reduced GC and PB compartments, indicating that antibody-mediated enhancement is T cell help dependent (Fig. 2 B and C). To examine whether antibody pre-infusion altered T cell responses, we employed Vav^Tg^Col1a^mCherry/+^ mice to measure early T cell activation and expansion and noted no significant differences between 3BNC117 IgG2a infused and control mice S3 A-C). To increase the sensitivity of our T cell assay we fused the Ovalbumin OVA^323-339^ peptide which is recognized by OT-II cells with high affinity to TM4-Core (Fig. S3 D-F). Transfer of OT-II cells labeled with the cell division tracking dye cell trace violet (CTV) into TM4-Core-OVA^323-339^ immunized mice revealed an equivalent extent of OT-II division between antibody infused mice and controls at both early (day 0-4) and later (day 4-7) time intervals (Fig. S3 G-L). Finally, despite the larger GCs, we also found no significant differences in T follicular helper (Tfh) cell numbers on day 7 after immunization M-O). Thus, although T cell help is essential for antibody-mediated enhancement of GC and PB responses, we found no significant antibody-dependent changes in the T cell compartment.

**Figure 2:**
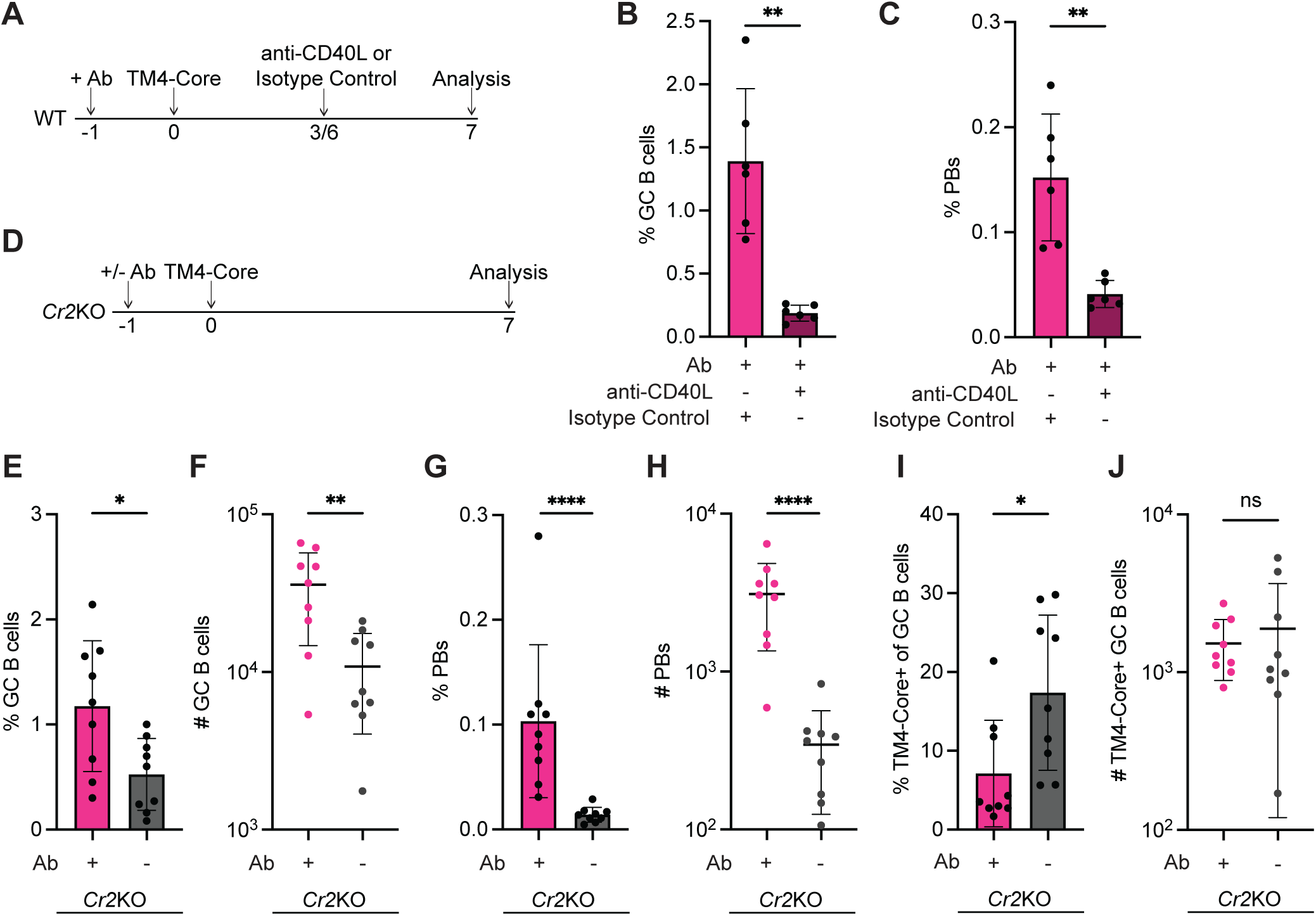
Antibody-mediated effects require CD40L but not CR1/2. **(A)** Experimental schematic for **B-C**. **(B)** Plot shows the percentage of GC B cells of B220+ cells in mice pre-infused with 3BNC117 IgG2a in concert with anti-CD40L (maroon, for **B-C**) or isotype control (pink, for **B-C**). **(C)** Plot shows the percentage of PBs in mice pre-infused with 3BNC117 IgG2a in concert with anti-CD40L or isotype control. **(D)** Experimental schematic for **E-J**. **(E)** Plot shows the percentage of GC B cells of B220+ cells in *Cr2*KO mice pre-infused with 3BNC117 IgG2a (+Ab, pink, for **E-J**) or no antibody (control, -Ab, grey, for **E-J**). **(F)** Plot shows the number of GC B cells in *Cr2*KO mice pre-infused with 3BNC117 IgG2a or control. **(G)** Plot shows the percentage of PBs in *Cr2*KO mice pre-infused with 3BNC117 IgG2a or control. **(H)** Plot shows the number of PBs in *Cr2*KO mice pre-infused with 3BNC117 IgG2a or control. **(I)** Plot shows the percentage of TM4-Core+ cells of GC B cells in *Cr2*KO mice pre- infused with 3BNC117 IgG2a or control. **(J)** Plot shows the number of TM4-Core+ GC B cells in *Cr2*KO mice pre-infused with 3BNC117 IgG2a or control. Data is displayed as mean ± SD. Each point represents a single mouse from at least two independent experiments. Statistical significance for **B-C** was determined using two-tailed unpaired t tests with Welch’s correction. Statistical significance for **E-J** was determined using two-tailed Mann-Whitney U tests. ns denotes non-significant, * denotes p ≤ 0.05, ** denotes p ≤ 0.01 and **** denotes p ≤ 0.0001.

### Complement receptors

Murine IgG2a is known to be an efficient activator of the complement cascade (Klaus et al., 1979; Neuberger and Rajewsky, 1981). To determine whether CR1/2 is required for enhanced GC and PB responses and the reduced fraction of detectable antigen-binding cells after antibody infusion, we employed *Cr2*KO mice which lack CR1/2 on all hematopoietic cells and stromal cells. Notably, 3BNC117 IgG2a infusion enhanced the magnitude of the GC and PB compartments in *Cr2*KO mice, and the GCs in antibody infused mice showed a reduction in the fraction of antigen-binding cells (Fig. 2D-J). These results indicate that CR1/2 is largely dispensable for antibody-mediated enhancement.

### FcγRs in GC and PB enhancement

To examine the role of type I FcγRs in antibody-mediated enhancement we employed FcRα null mice which lack FcγRI, FcγRIIb, FcγRIII and FcγRIV but retain expression of FcRn, the neonatal Fc receptor that mediates antibody recycling, and type II Fc receptors, CD209 and CD23 (Pincetic et al., 2014; Smith et al., 2012). GC and PB responses to TM4-Core in 3BNC117 IgG2a infused FcRα null mice were similar to controls, indicating that FcγRs are essential for enhancement (Fig. 3 A-E). To confirm that FcγRs are necessary, we repeated the experiment in WT mice using 3BNC117 -IgG2a, -IgG1, or -IgG1 D265A (Fig. 3 A). Murine IgG1 binds selectively to FcγRIIb, the low affinity inhibitory receptor, and to FcγRIII, a low affinity activating receptor. The D265A mutation ablates FcγR binding (Bournazos et al., 2014; Nimmerjahn et al., 2005; Nimmerjahn and Ravetch, 2005). To control for antigen specificity, we used GO53 IgG2a, an antibody that does not bind to TM4-Core (Wardemann et al., 2003). Consistent with the results obtained with FcRα null mice, 3BNC117 IgG1 D265A infusion was not significantly different from controls (Fig. 3 G and H). In contrast, 3BNC117 IgG1 showed an intermediate phenotype (Fig. 3 G and H). We conclude that type I FcγRs are essential for antibody-mediated enhancement of GC and PB responses.

**Figure 3:**
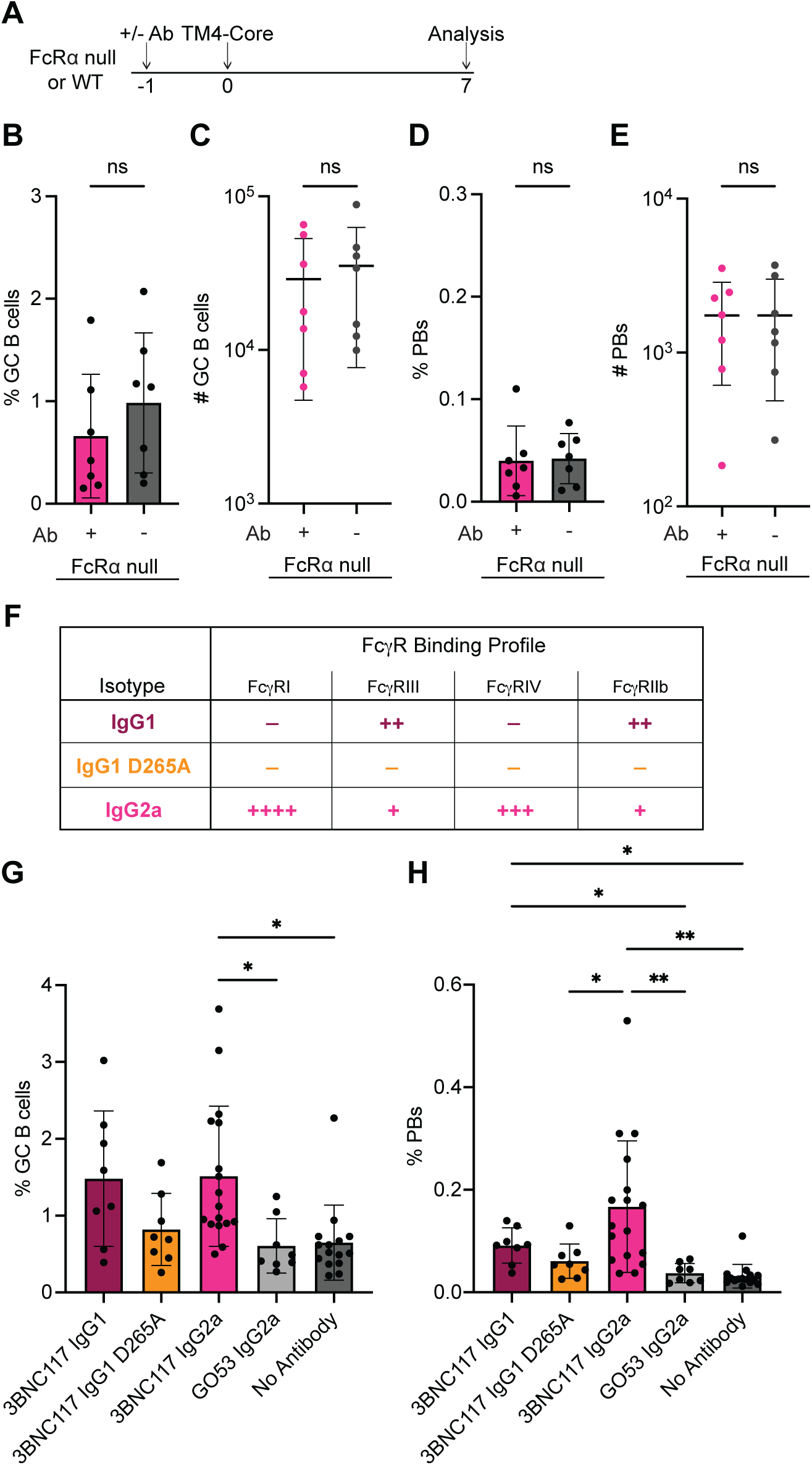
Enhanced magnitude of GC and PB compartments is dependent on FcγRs. **(A)** Experimental schematic for **B-E** and **G-H**. **(B)** Plot shows the percentage of GC B cells of B220+ cells in FcRα null mice pre-infused with 3BNC117 IgG2a (+Ab, pink, for **B-E**) or no antibody (control, -Ab, grey, for **B-E**). **(C)** Plot shows the number of GC B cells in FcRα null mice pre-infused with 3BNC117 IgG2a or control. **(D)** Plot shows the percentage of PBs in FcRα null mice pre-infused with 3BNC117 IgG2a or control. **(E)** Plot shows the number of PBs in FcRα null mice pre-infused with 3BNC117 IgG2a or control. **(F)** Chart shows murine isotypes and their respective murine FcγR binding profiles. Chart adapted from Bournazos et al., *Cell*, 2014 (Bournazos et al., 2014). **(G)** Plot shows the percentage of GC B cells of B220+ cells in mice pre-infused with 3BNC117 IgG1 (maroon, for **G-H**), 3BNC117 IgG1 D265A (orange, for **G-H**), 3BNC117 IgG2a (pink, for **G-H**), GO53 IgG2a (light grey, for **G-H**) or no antibody (control, dark grey, for **G-H**). **(H)** Plot shows the percentage of PBs of in mice pre-infused with 3BNC117 IgG1, 3BNC117 IgG1 D265A, 3BNC117 IgG2a, GO53 IgG2a or control. Data is displayed as mean ± SD. Each point represents a single mouse from at least two independent experiments. Statistical significance for **B-E** was determined using two-tailed unpaired t tests. Statistical significance for **G-H** was determined using Brown-Forsythe and Welch ANOVA tests with multiple comparisons. ns denotes non-significant, * denotes p ≤ 0.05 and ** denotes p ≤ 0.01.

### Antigen-specific cells

In addition to increasing the magnitude of GC and PB responses, pre-existing antibody reduced the fraction of detectable antigen-binding cells within GCs. To examine whether this phenomenon is dependent on FcγRs, we determined the fraction of antigen-binding cells in the GC on day 7 after immunization with TM4-Core in FcRα null mice 4 A). FcRα null mice infused with antibody produced GCs with a reduced fraction of antigen-binding cells compared to controls (Fig. 4 B and C). Consistent with this finding, and in contrast to their effects on GC and PB magnitude, pre-infusion of 3BNC117 -IgG1, -D265A, or -IgG2a all reduced the fraction of antigen-binding cells in the GC (Fig. 4 D). Thus, antibodies mediate GC diversification in an antigen-specific but FcγR independent manner.

**Figure 4:**
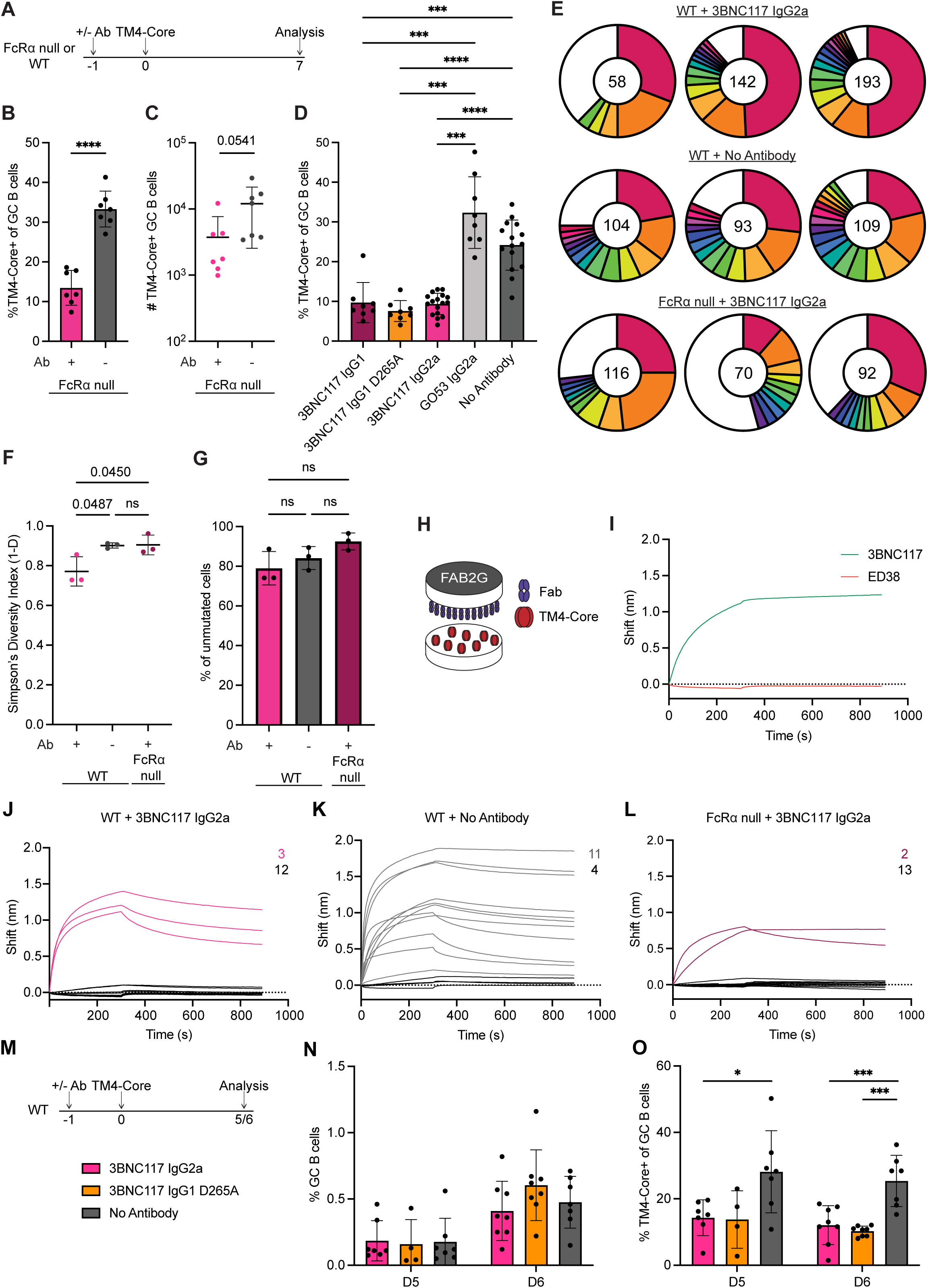
Antibody infusion elicits GC and PB compartments with reduced antigen-binding by an FcγR independent mechanism. **(A)** Experimental schematic for **B-L**. **(B)** Plot shows the percentage of TM4-Core+ cells of GC B cells in FcRα null mice pre-infused with 3BNC117 IgG2a (+Ab, pink, for **B-C**) or no antibody (control, -Ab, grey, for **B-C**). **(C)** Plot shows the number of TM4-Core+ GC B cells in FcRα null mice pre-infused with 3BNC117 IgG2a or control. **(D)** Plot shows the percentage of TM4-Core+ cells of GC B cells in WT mice pre-infused with 3BNC117 IgG1 (maroon), 3BNC117 IgG1 D265A (orange), 3BNC117 IgG2a (pink), GO53 IgG2a (light grey) or no antibody (control, dark grey). **(E)** Pie charts show clonal distribution of sequenced immunoglobulin genes from PBs isolated from WT mice pre-infused with 3BNC117 IgG2a (top), WT mice pre-infused with no antibody (middle) and FcRα null mice pre-infused with 3BNC117 IgG2a (bottom). Number in the middle represents the total sequences obtained; colored slices represent expanded clones and white slices represent singlets. **(F)** Plot shows Simpson’s Diversity Index (1-D) of PBs isolated from WT mice pre-infused with 3BNC117 IgG2a (pink, for **F-G**), WT mice pre-infused with no antibody (grey, for **F-G**) or FcRα null mice pre-infused with 3BNC117 IgG2a (maroon, for **F-G**). **(G)** Plot shows the percentage of unmutated cells from WT mice pre-infused with 3BNC117 IgG2a, WT mice pre-infused with no antibody or FcRα null mice pre-infused with 3BNC117 IgG2a. **(H)** Schematic of BLI set-up. Cloned Fabs from PBs are immobilized on a FAB2G sensor and immersed in a solution of TM4-Core. **(I)** Graph shows BLI traces for positive control Fab 3BNC117 (green) and negative control Fab ED38 (red). **(J)** Graph shows BLI traces for Fabs produced from BCR sequences of PBs isolated from WT mice pre-infused with 3BNC117 IgG2a. Binders and number of binders are denoted in pink traces/text. Non-binders and number of non-binders are denoted in black traces/text. **(K)** Graph shows BLI traces for Fabs produced from BCR sequences of PBs isolated from control WT mice pre-infused with no antibody. Binders and number of binders are denoted in grey traces/text. Non-binders and number of non-binders are denoted in black traces/text**. (L)** Graph shows BLI traces for Fabs produced from BCR sequences of PBs isolated from FcRα null mice pre-infused with 3BNC117 IgG2a. Binders and number of binders are denoted in maroon traces/text. Non-binders and number of non-binders are denoted in black traces/text**. (M)** Experimental schematic for **N-O** with legend. **(N)** Graph shows the percentage of GC B cells of B220+ cells on day 5 (D5) and day 6 (D6) post immunization in mice pre-infused with 3BNC117 IgG2a (pink, for **N-O**), 3BNC117 IgG1 D265A (orange, for **N-O**) or no antibody (control, grey, for **N-O**). **(O)** Graph shows the percentage of TM4-Core+ cells of GC B cells on day 5 (D5) and day 6 (D6) post immunization in mice pre-infused with 3BNC117 IgG2a, 3BNC117 IgG1 D265A or control. Data is displayed as mean ± SD. For **B-D** and **N-O**, each dot represents a single mouse from at least two independent experiments (only one experiment for 3BNC117 IgG1 D265A on day 5 in **N-O**). For **F-G**, each dot represents the results from sequencing data of a single mouse. Statistical significance for **B-C** was determined using two-tailed unpaired t tests. Statistical significance for **D** was determined using a Brown-Forsythe and Welch ANOVA test with multiple comparisons. Statistical significance for **F-G** and **N-O** was determined using an ordinary one-way ANOVA with multiple comparisons (**N-O** was done with separate ANOVAs for each timepoint). ns denotes non-significant, * denotes p ≤ 0.05, *** denotes p ≤ 0.001 and **** denotes p ≤ 0.0001.

Plasmablasts express low levels of surface immunoglobulin making flow-cytometric detection of antigen binding less reliable (Pelletier et al., 2006; Ellyard et al., 2004). To examine the effects of antibody pre-infusion on the composition of the PB compartment, single PBs were purified and their immunoglobulin genes were sequenced 7 days after immunization with TM4-Core (Fig. 4 A). At this early stage in the immune response, PBs were clonally expanded and diverse with little to no somatic hypermutation (Fig. 4 E-G). To determine whether there were qualitative differences in the antibodies produced by PBs obtained from WT or FcRα null mice pre-infused with 3BNC117 IgG2a or no antibody, we expressed Fabs representing the 5 most expanded clones in each of 9 mice and tested them for binding to TM4-Core by bio-layer interferometry (BLI) H and I). Whereas 11/15 Fabs from WT controls bound to antigen, only 3/15 of Fabs from 3BNC117 IgG2a infused WT and 2/15 Fabs from FcRα null mice did so (Fig. 4 J-L). Thus, while the antibody-mediated increase in the magnitude of the GC and PB response is FcγR dependent, the parallel reduction in the fraction of detectable antigen-binding cells is FcγR independent.

To understand whether the FcγR independent reduction in detectable antigen-binding cells is a result of differential GC seeding, we examined responses 5 and 6 days after immunization (Fig. 4 M). Even at this early time point when GCs are first developing, both 3BNC117 IgG2a and IgG1 D265A pre-infusion reduced the fraction of antigen-binding cells in the GC (Fig. 4 N and O). We conclude that antibodies can influence the diversity of B cells in early GCs by altering recruitment in an FcγR independent manner.

### FcγRs in immune complex retention

To examine how FcγRs may facilitate larger magnitude GC and PB responses, we investigated the distribution of immune complexes (ICs) in the LNs of WT and FcRα null mice. To facilitate IC detection, we used human 3BNC117 IgG1 which exhibits a comparable binding profile to murine FcγRs as murine IgG2a (Dekkers et al., 2017). Moreover, human 3BNC117 IgG1 pre-infusion produces similar effects on the GC and PB compartments as murine IgG2a (Fig. S4 A-D).

WT and FcRα null mice were pre-infused with human 3BNC117 IgG1 and subsequently immunized with TM4-Core (Fig. 5 A). Immunofluorescence staining of draining LNs in WT mice showed ICs on follicular dendritic cells (FDCs) on day 1 which persisted at nearly the same level until day 4 after immunization (Fig. 5 B and C). ICs were also detectable in FcRα null mice on day 1 but were significantly reduced by day 4 compared to WT (Fig. 5 B and C). In addition to FDCs, FcγR dependent IC retention was found on the surface of F4/80+ macrophages, migratory dendritic cells and in the corticomedullary region of the draining LN, where PBs accumulate (Fig. 5 B and D; E-J). We conclude that FcγRs promote enhanced IC retention in GCs and the PB-rich corticomedullary regions.

**Figure 5:**
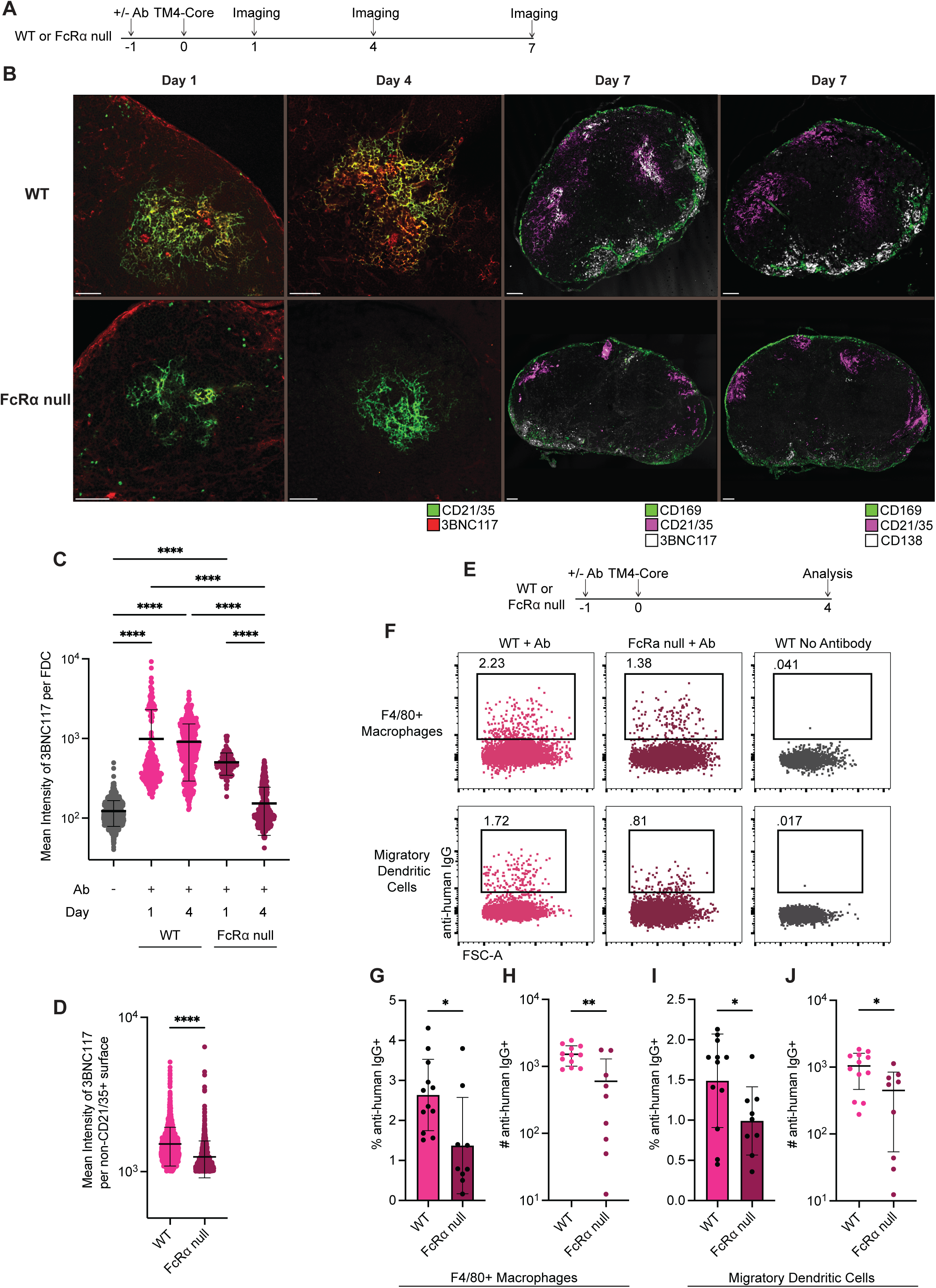
Antibody mediates enhanced IC retention in an FcγR dependent manner on FDCs and in the lymph node medulla. **(A)** Experimental schematic for **B-D**. **(B)** Confocal microscopy images of popliteal LNs from WT or FcRα null mice at the timepoints indicated according to the above schematic. For the left two columns of images at day 1 and day 4, sections are stained with anti-CD21/35 (green) and anti-human IgG for 3BNC117 (red). Scale bars represent 50 µM. For the third column, images show day 7 slices stained with anti-human IgG for 3BNC117 (white), anti-CD21/35 (pink) and anti-CD169 (green). Scale bars represent 100 µM. For the fourth column, images show day 7 slices stained with anti-CD138 (white), anti-CD21/35 (pink) and anti-CD169 (green). Scale bars represent 100 µM. **(C)** Quantification of the mean intensity of infused human 3BNC117 IgG1 antibody detected per FDC surface using anti-human IgG. Each dot represents the mean intensity of antibody detected on a single surface, with dots pooled from at least 2 mice. Grey represents WT mice pre-infused with no antibody (control), pink represents WT mice pre-infused with human 3BNC117 IgG1 and maroon represents FcRα null mice pre- infused with human 3BNC117 IgG1. **(D)** Quantification of the mean intensity of infused human 3BNC117 IgG1 antibody detected on surfaces other than those colocalized with CD21/35 expression using anti-human IgG. Each dot represents the mean intensity of antibody on a single detected surface, with dots pooled from at least 2 mice. Pink represents WT mice pre-infused with human 3BNC117 IgG1 and maroon represents FcRα null mice pre-infused with human 3BNC117 IgG1. **(E)** Experimental schematic for **F-J**. **(F)** Representative flow cytometry plots for extracellular detection of human 3BNC117 IgG1 using anti-human IgG. Top shows gating for detection of human 3BNC117 IgG1 on F4/80+ macrophages (B220-, F4/80+); bottom shows gating for detection of human 3BNC117 IgG1 on migratory dendritic cells (B220-, F4/80-, CD169-, MHC-II^high^, CD11c^int^). Flow plots for WT mice infused with human 3BNC117 IgG1 (left panels, pink), FcRα null mice infused with human 3BNC117 IgG1 (middle panels, maroon) and WT mice pre-infused with no antibody (control, right panels, grey). **(G)** Plot shows the percentage of F4/80+ macrophages with detectable human 3BNC117 IgG1 using anti-human IgG in WT (pink, for **G-J**) and FcRα null (maroon, for **G-J**) mice. **(H)** Plot shows the number of F4/80+ macrophages with detectable human 3BNC117 IgG1 using anti-human IgG in WT and FcRα null mice. **(I)** Plot shows the percentage of migratory dendritic cells with detectable human 3BNC117 IgG1 using anti-human IgG in WT and FcRα null mice. **(J)** Plot shows the number of migratory dendritic cells with detectable human 3BNC117 IgG1 using anti-human IgG in WT and FcRα null mice. Data is displayed as mean ± SD. Statistical significance for **C** was determined using a Kruskal-Wallis test with multiple comparisons. Statistical significance for **D** was determined using a two-tailed Mann-Whitney U test. Statistical significance for **G-J** was determined using two-tailed unpaired t tests. * denotes p ≤ 0.05, ** denotes p ≤ 0.01 and **** denotes p ≤ 0.0001.

## Discussion

Understanding how antibodies modulate immune responses is of the utmost importance to optimizing vaccination regimens. For example, pre-infusion of two monoclonal antibodies into humans 90 days before SARS-CoV-2 vaccination altered both the nature of the immunodominant epitopes targeted and the affinity of antibodies elicited by the vaccine (Schaefer-Babajew et al., 2023). Here, we examined the mechanisms by which antibodies modulate the magnitude and quality of immune responses.

We find that in WT mice pre-existing high affinity antigen-specific antibodies increase the magnitude of both the GC and PB compartment in a FcγR dependent manner. Murine IgG2a used in most of our experiments is similar to human IgG1 which is the dominant isotype used for defense against viral pathogens and binds to all activating and inhibitory FcγRs (Bournazos and Ravetch, 2017; Bruhns et al., 2009; Dekkers et al., 2017). Our findings with virus-derived protein antigens aligns with and extends earlier work that was largely limited to hapten immunogens and measurements of serologic responses, wherein antibody-mediated modulation of immune responses was also found to be FcγR dependent (Ståhl and Heyman, 2001; Getahun et al., 2004; Wernersson et al., 1999).

FcγRIIb is the only FcγR expressed on FDCs and IC retention on FDCs is essential for GC responses (Radoux et al., 1985; Qin et al., 2000; Yoshida et al., 1993; Poel et al., 2019; Wang et al., 2011). However, these cells also express CR1/2 that can play a dominant role in IC retention (Heesters et al., 2013; Fang et al., 1998; Martínez-Riaño et al., 2023; Phan et al., 2007). The finding that ablation of FcγRs significantly reduces IC retention on FDCs suggests that in the case of antibody isotypes with broad FcγR binding, FcγRs are essential for optimal IC trapping. Moreover, the observation of FcγR dependent IC retention on hematopoietic cells in the LN such as macrophages and dendritic cells indicates that these receptors can promote antigen retention on the surface of cells other than FDCs. The anatomical colocalization of plasma cells and ICs in the corticomedullary region reinforces the idea that increased antigen retention can increase the overall magnitude of immune responses, a concept aligned with the enhanced GC, T cell and serum Ig responses observed in escalating dose regimens (Tam et al., 2016). Finally, in addition to antigen trapping, ICs can inhibit B cell receptor signaling through FcγRIIb and also enhance antigen uptake for processing and presentation as peptide-MHC to cognate T cells (Muta et al., 1994; Dvorscek et al., 2024; Kalergis and Ravetch, 2002; Tang et al., 2021).

Our experiments illustrate that pre-existing antibody can also alter the quality of the GC and PB compartments by reducing the fraction of detectable antigen-binding cells. This mechanism was independent of FcγR binding and was documented at the earliest stages of the response, suggesting that ICs increase the apparent valency of the antigen and thereby lower the threshold for B cell recruitment. Our experiments are in agreement with and extend the observations in transgenic mice and humans showing that pre- existing antibody can recruit low affinity B cells into the immune response (Dvorscek et al., 2024; Schaefer-Babajew et al., 2023; Hägglöf et al., 2023). However, these effects appear to be a function of valency and not specific for ICs. Increasing antigen valency from a HIV-1 tetramer to a 60-mer was sufficient to increase the size of the GC, and decrease the fraction of antigen-binding cells, while also conserving the absolute numbers of antigen-binding cells in the GC (Kato et al., 2020).

In conclusion, antibodies play a critical role in immune memory beyond direct pathogen clearance or neutralization. Pre-existing antibodies can mask immunodominant epitopes and diversify immune responses by shifting the focus to subdominant epitopes (Tas et al., 2022; Dvorscek et al., 2024; McNamara et al., 2020; Schaefer-Babajew et al., 2023). Additionally, ICs diversify immune response by lowering the threshold for B cell recruitment in an FcγR independent manner. Finally, ICs enhance the magnitude of the response and trap antigens in lymphoid organs via an FcγR dependent mechanism, thereby prolonging their interaction with cellular components of the adaptive immune system.

### Author contributions

M.C., A.J.M. and M.C.N. conceived, designed and analyzed all experiments. A.G., B.H., and L.S. provided critical reagents. G.S.S.S. performed bioinformatic analysis. H.H., J.M., S.B., and J.R. provided critical tools and guidance to this study. M.C. and M.C.N. wrote the manuscript with input from all authors.

## Acknowledgements

We thank T. Eisenreich and K. Yao for technical help and mouse colony management, K. Gordon for performing cell sorting and M. Jankovic and T. Waldetario for essential laboratory support. We thank the Ravetch laboratory for providing critical mouse models used in this work, and all members of the Nussenzweig laboratory for helpful discussion. Confocal microscopy was performed at the Rockefeller University’s Bio-Imaging Resource Center.

## Online supplemental material

**Fig. S1** shows the gating strategy for GC B cells, PBs and TM4-Core+ GC B cells. **Fig. S2** shows antibody modulation with an alternative antibody-antigen combination. **Fig. S3** shows analysis of antibody pre-infusion on various T cell compartments. **Fig. S4** shows antibody modulation in mice using human immunoglobulins as opposed to murine immunoglobulins.

## Data availability

Data supporting the findings of this work are available in the article and supplementary materials.

---

**Supplemental Figure 1:**
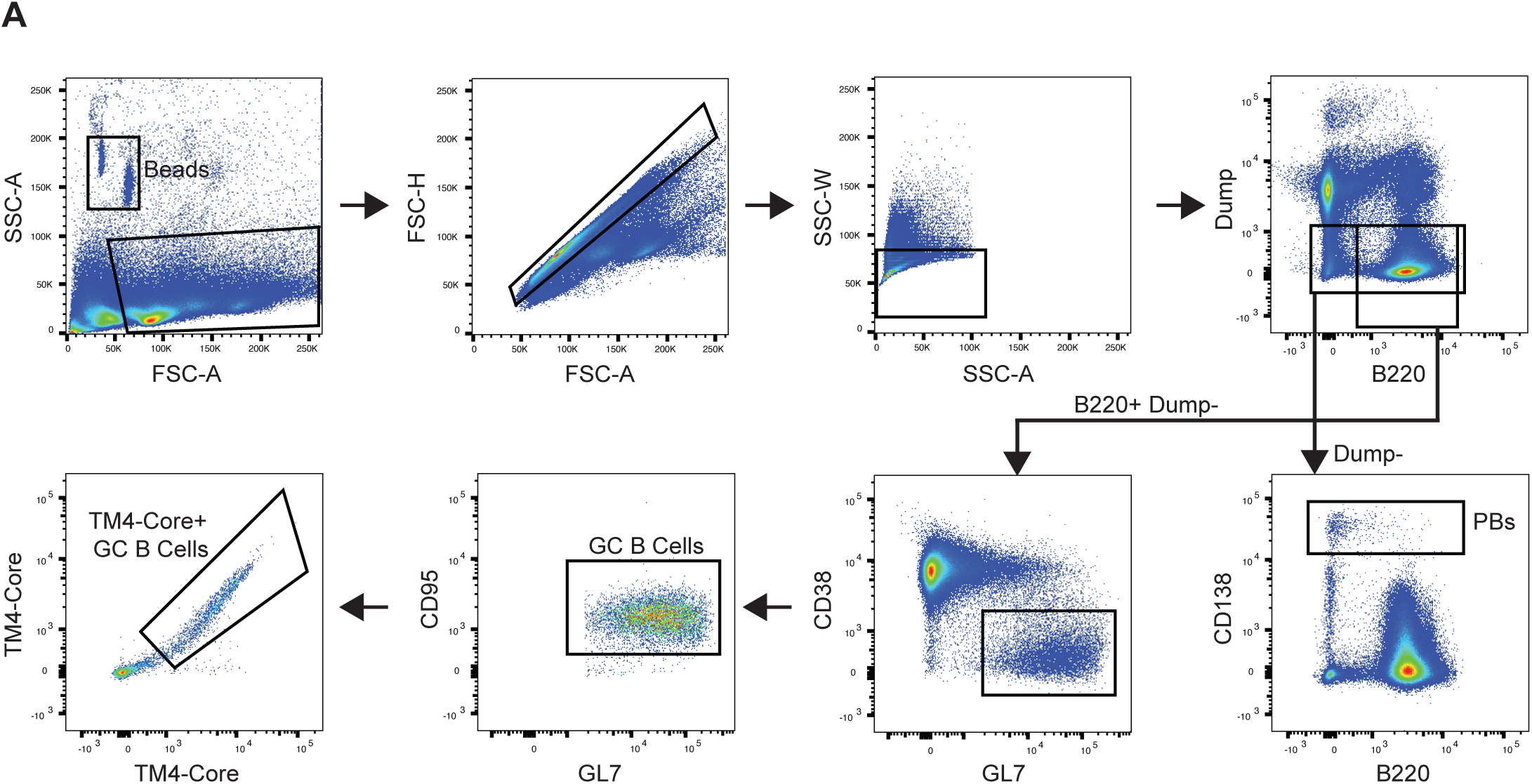
Gating strategy for GC, PB and TM4-Core+ GC B cell compartments. **(A)** Gating strategy for GC B cells (Dump-, B220+, CD38-, GL7+, CD95+), PBs (Dump-, CD138+) and TM4-Core antigen-bait+ GC B cells (Dump-, B220+, CD38-, GL7+, CD95+, TM4-Core antigen-bait+). Dump channel includes Live/Dead, CD4, CD8a, NK1.1 and F4/80.

**Supplemental Figure 2:**
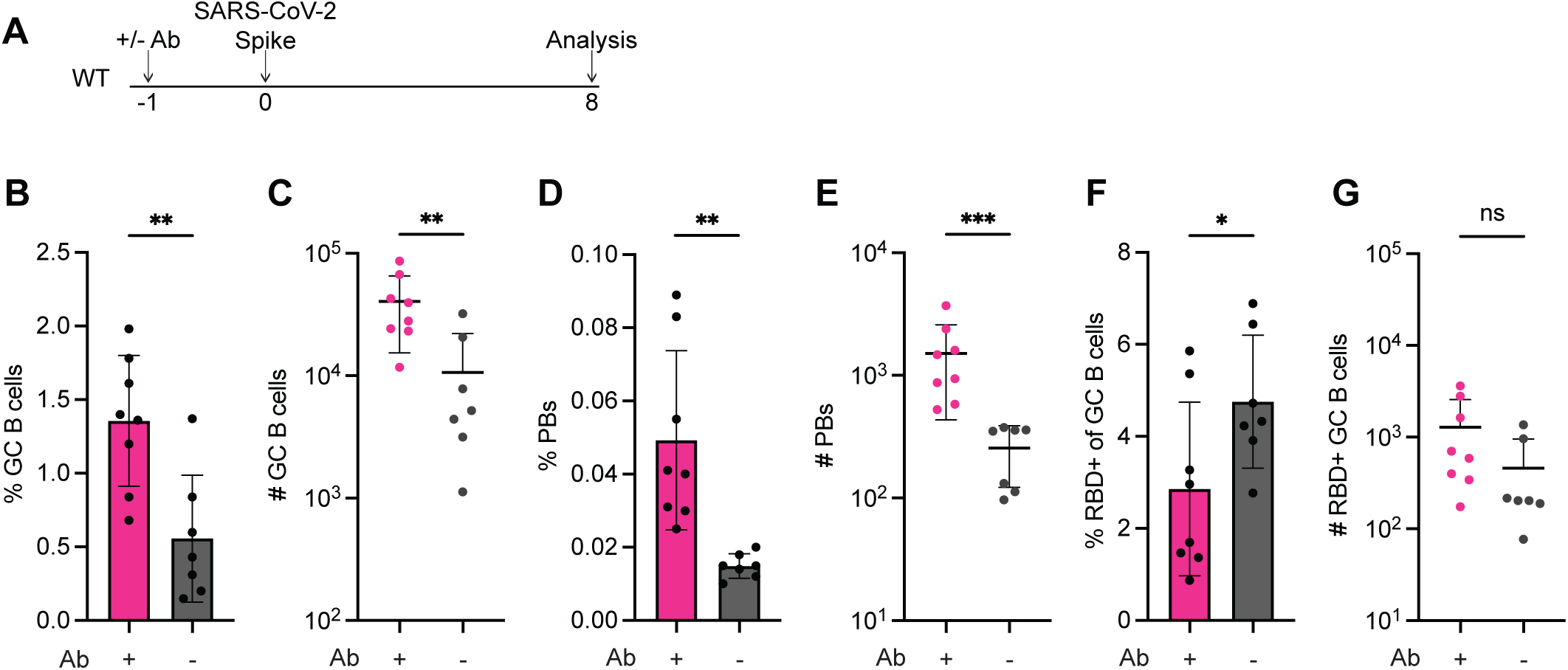
Pre-infusion with C144 IgG2a prior to Spike immunization similarly modulates the immune response. **(A)** Experimental schematic for **B-G**. **(B)** Plot shows the percentage of GC B cells of B220+ cells in mice pre-infused with C144 IgG2a (+Ab, pink, for **B-G**) or no antibody (control, -Ab, grey, for **B-G**). **(C)** Plot shows the number of GC B cells in mice pre-infused with C144 IgG2a or control. **(D)** Plot shows the percentage of PBs in mice pre-infused with C144 IgG2a or control. **(E)** Plot shows the number of PBs in mice pre-infused with C144 IgG2a or control. **(F)** Plot shows the percentage of RBD+ cells of GC B cells in mice pre-infused with C144 IgG2a or control. **(G)** Plot shows the number of RBD+ GC B cells in mice pre-infused with C144 IgG2a or control. Data is displayed as mean ± SD. For **B-G** each dot represents a single mouse from at least two independent experiments. Statistical significance for **B**, **D** and **F** was determined using two-tailed unpaired t tests. Statistical significance for **C**, **E** and **G** was determined using two-tailed Mann-Whitney U tests. ns denotes non-significant, * denotes p ≤ 0.05, ** denotes p ≤ 0.01 and *** denotes p ≤ 0.001.

**Supplemental Figure 3:**
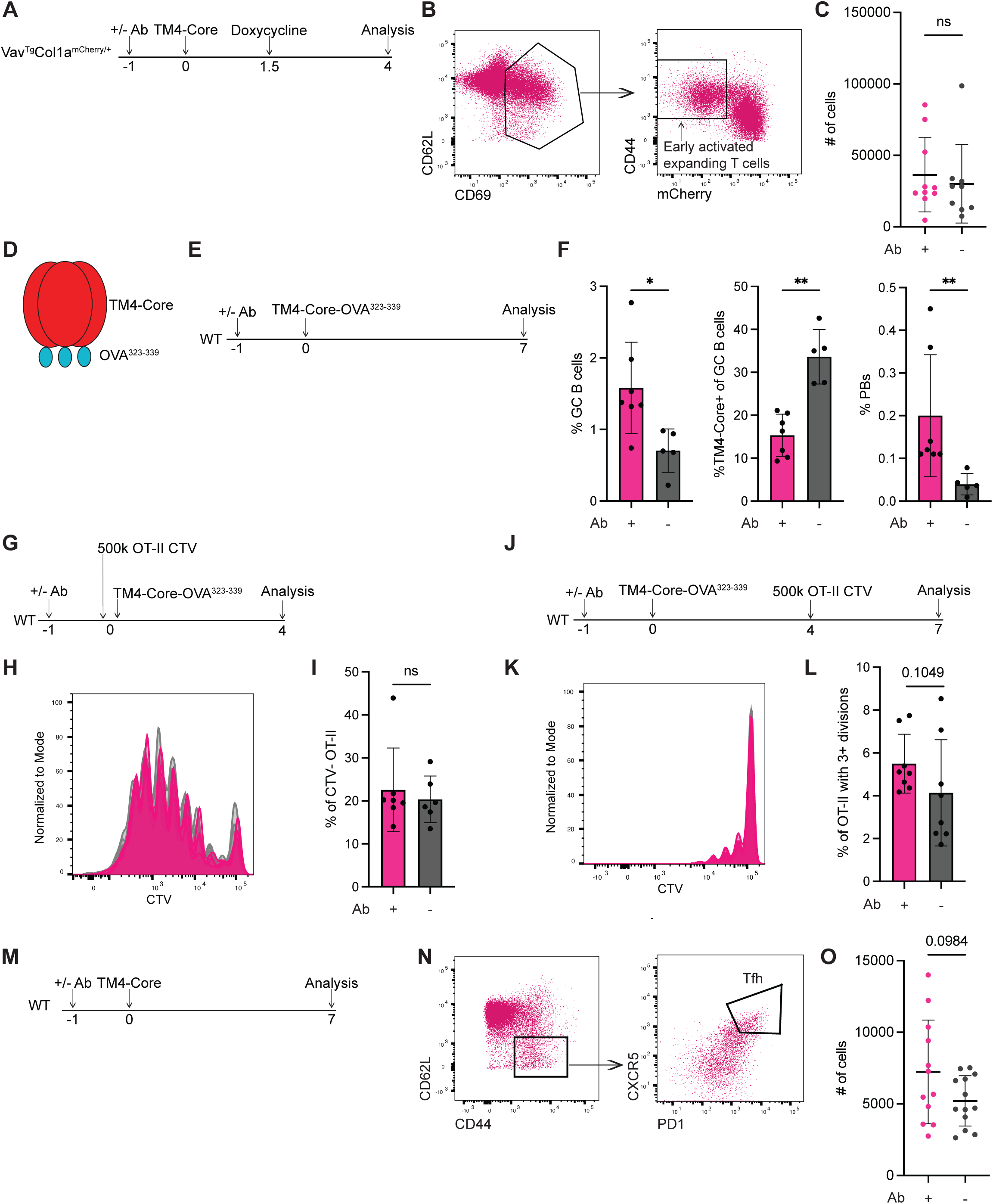
Antibody does not detectably influence T cell compartments. **(A)** Experimental schematic for **B-C**. **(B)** Representative flow cytometry plots of gating strategy for early activated expanding T cells (CD4+, CD62L CD69 transitional, CD44^high^, mCherry^low^). **(C)** Plot showing the number of early activated expanding T cells in mice pre-infused with 3BNC117 IgG2a (+Ab, pink) or no antibody (control, -Ab, grey). **(D)** Representation of TM4-Core-OVA^323-339^ immunogen. **(E)** Experimental schematic for **F**. **(F)** Plot shows the percentage of GC B cells of B220+ cells (left), the percentage of TM4-Core+ cells of GC B cells (middle) and the percentage of PBs in mice pre-infused with 3BNC117 IgG2a (+Ab, pink) or no antibody (control, -Ab, grey). **(G)** Experimental schematic for **H-I**. **(H)** Representative histogram overlays of OT-II CTV dilution normalized to the mode for mice pre-infused with 3BNC117 IgG2a antibody (pink) or no antibody (grey). **(I)** Plot shows the percentage of fully CTV diluted OT-II cells amongst all OT-II cells in mice pre-infused with 3BNC117 IgG2a (+Ab, pink) or no antibody (control, -Ab, grey). **(J)** Experimental schematic for **K-L**. **(K)** Representative histogram overlays of OT-II CTV dilution normalized to the mode for mice pre-infused with 3BNC117 IgG2a (pink) or no antibody (grey). **(L)** Plot shows the percentage of OT-II cells with 3 or more divisions amongst all OT-II cells for mice pre-infused with 3BNC117 IgG2a (+Ab, pink) or no antibody (control, -Ab, grey). **(M)** Experimental schematic for **N-O**. **(N)** Representative flow cytometry plots of gating strategy for Tfh cells (CD4+, CD62L-, CD44+, PD1^high^, CXCR5^high^). **(O)** Plot showing the absolute number of Tfh cells in mice pre-infused with 3BNC117 IgG2a (+Ab, pink) or no antibody (control, -Ab, grey). Data is displayed as mean ± SD. For **C**, **F**, **I**, **L** and **O**, each dot represents a single mouse from at least two independent experiments. Statistical significance for **C**, **F I**, and **L** was determined using two-tailed Mann-Whitney U tests. Statistical significance for **O** was determined using a two-tailed unpaired t test with Welch’s correction. ns denotes non-significant, * denotes p ≤ 0.05 and ** denotes p ≤ 0.01.

**Supplemental Figure 4:**
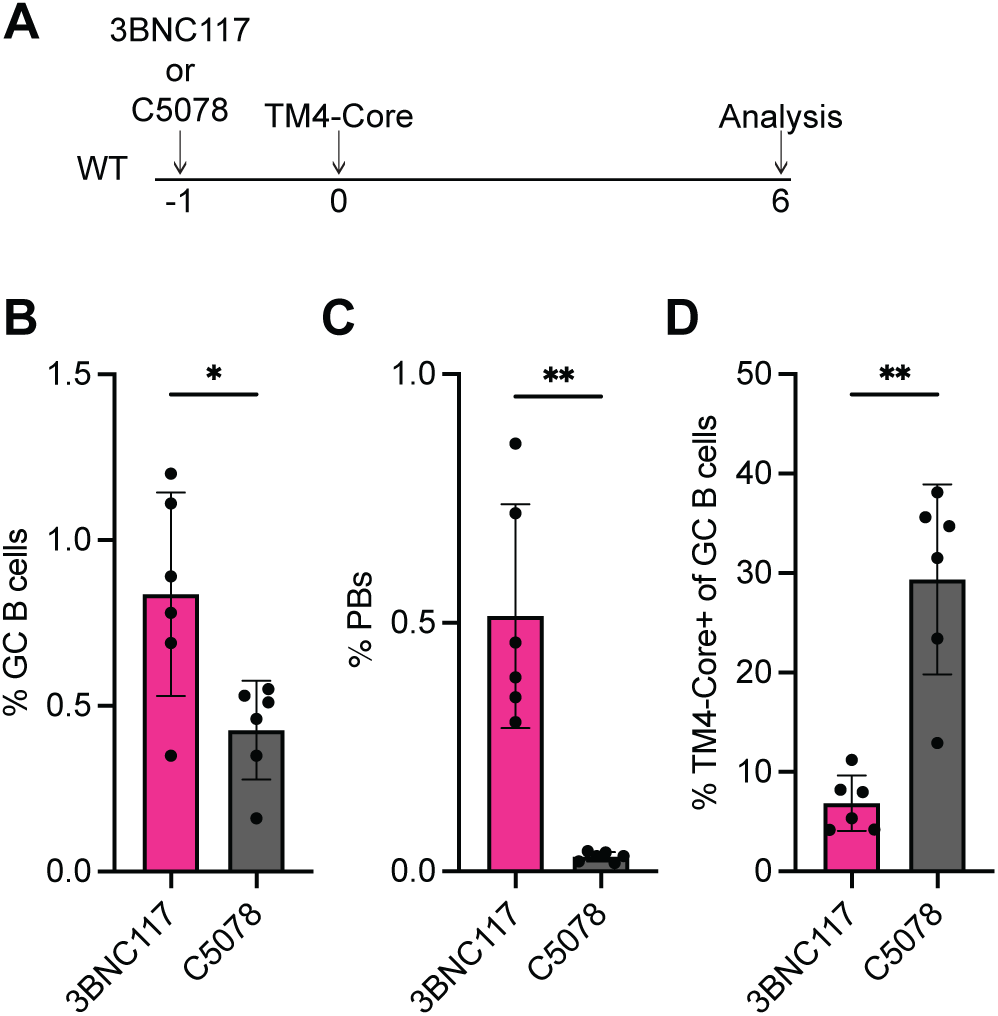
Human IgG1 infusion in mice produces similar modulation phenomenon to murine IgG2a infusion. **(A)** Experimental schematic for **B-D**. **(B)** Plot shows the percentage of GC B cells of B220+ cells in mice pre-infused with human 3BNC117 IgG1 (pink, for **B-D**) or human C5078 IgG1 antibody (control, grey, for **B-D**). **(C)** Plot shows the percentage of PBs in mice pre-infused with human 3BNC117 IgG1 or control. **(D)** Plot shows the percentage of TM4-Core+ cells of GC B cells in mice pre-infused with human 3BNC117 IgG1 or control. Data is displayed as mean ± SD. For **B-D** each dot represents a single mouse from two independent experiments. Statistical significance for **B-D** was determined using two-tailed unpaired t tests with Welch’s correction. * denotes p ≤ 0.05 and ** denotes p ≤ 0.01.

## Materials & Methods

### Mice

All mice were housed at Rockefeller University at 22°C on a 12 hour dark/light cycle with access to food and water. WT C57BL/6J (Jax: 000664), WT CD45.1 B6 (Jax: 002014) and *Cr2*KO (Jax: 008225) mice were purchased from Jackson Laboratories (Molina et al., 1996; Shen et al., 1985). FcRα null mice were kindly provided by Dr. Jeffrey Ravetch and Dr. Stylianos Bournazos (Smith et al., 2012). OT-II were bred and maintained at Rockefeller. Vav^Tg^Col1a^mCherry/+^ mice were generated in house and previously described (Gitlin et al., 2014). Male and female mice between the ages of 6-12 weeks were used for experiments. All procedures were performed following protocols approved by the IACUC.

### Immunizations

TM4-Core, TM4-Core-OVA^323-339^ and Spike immunizations were performed via footpad injection. 5 µg of each immunogen was resuspended at a 2:1 ratio of immunogen in 1x PBS to Imject Alum (Thermo Scientific 77161) in a total volume of 25 µL/footpad.

### Antibody infusions

100 µg of a given antibody was resuspended in 100-150 µL of 1x PBS and infused via retro-orbital injection. For most cases, control mice receiving no antibody were infused with 100-150 µL of 1x PBS.

### Doxycycline and anti-CD40L treatment

For anti-CD40L treatment, mice intravenously received either two 300 µg injections of anti-CD40L (Clone: MR1, BioXCell BE0017-1) or an armenian hamster IgG isotype control (BioXCell BE0091) in 100 µL of 1x PBS. For doxycycline, mice received 2 mg of doxycycline hyclate (Sigma-Aldrich D9891) in 200 µL of 1x PBS via intraperitoneal injection on the day indicated. To maintain suppression of mCherry throughout the remainder of the experiment, mice received 500 mL water bottles containing 2.5 g of sucrose (Sigma-Aldrich S0389) and 0.5 g of doxycycline.

### Immunogen and antigen-bait production

TM4-Core, TM4-Core gp120 and TM4-Core-OVA^323-339^ were provided by Dr. Leonidas Stamatatos or produced in house using a vector provided by Dr. Leonidas Stamatatos (McGuire et al., 2014; Dosenovic et al., 2018) . TM4-Core-OVA^323-339^ was generated by cloning the OVA^323-339^ peptide sequence into the carboxyl terminus of the protein. Spike and RBD protein were similarly produced in house. All immunogens and antigen-baits were produced via transient transfection of Expi293™ cells (Thermo Fisher A14527) with the Expifectamine™ 293 Expression System Kit (Thermo Fisher A14525) and collected 5-7 days post transfection for purification. TM4-Core and TM4-Core-OVA^323-339^ were purified using Galanthus Nivalis Lectin (Vector Laboratories AL-1243-5) and subsequently underwent size-exclusion chromatography to obtain their native trimeric form. Spike, RBD and TM4-Core gp120 were all purified using Ni Sepharose™ 6 Fast Flow (Millipore Sigma GE17-5318-03). Spike was also subjected to size-exclusion chromatography after purification.

### Antibody production

Most antibodies were produced in house via transient transfection of Expi293™ cells with the Expifectamine™ 293 Expression System Kit as aforementioned. Antibody vectors for murine IgG2a, IgG1 and IgG1 D265A were kindly provided by Dr. Jeffrey Ravetch and Dr. Stylianos Bournazos (Bournazos et al., 2014). Anti-CD40L (Clone: MR1, BioXCell BE0017-1) and the armenian hamster isotype control (BioXCell BE0091) were purchased from BioXCell. Murine IgG2a, IgG1, IgG1 D265A and human IgG1 were purified using Protein G Sepharose™ Fast Flow resin (Cytvia 17061806) and subsequently buffer exchanged into 1x PBS. For all antibodies, validation of production and purification was performed via gel electrophoresis. Briefly, antibodies were incubated with NuPAGE™ Sample Reducing Agent (Invitrogen NP0009) and NuPAGE™ LDS sample buffer (Invitrogen NP0007), heated shortly at 95°C and run on NuPAGE™ Bis-Tris Mini Protein Gels (Thermo Fisher NP0321Box). Protein gels were stained using One-Step Blue® Protein Gel Stain (Biotium 21003-1L) and subsequently imaged using an Amersham™ ImageQuant™ 800 (Cytvia 29399481). Antibodies that had higher than acceptable levels of LPS underwent LPS removal prior to injection using 10% Triton X-114 (Sigma-Aldrich X114).

### Fab production

Fabs were cloned using BCR sequences obtained from immunoglobulin sequencing of single PBs. eBlocks for heavy and light chain sequences were ordered from Integrated DNA Technologies (IDT) and underwent assembly into human Fab and human kappa expression vectors using NEBuilder® HiFi DNA Assembly Master Mix (New England Biolabs E2621). Fabs were similarly produced via transient transfection as antibodies above and purified using Ni Sepharose™ 6Fast Flow (Millipore Sigma GE17-5318-03). Fabs were buffer exchanged into 1x PBS and checked for production via gel electrophoresis as above.

### Antigen-bait generation

For TM4-Core and TM4-Core-OVA^323-339^ immunizations, biotinylated TM4-Core gp120, a monomeric version of TM4-Core lacking the trimerization domain, was employed as flow cytometric antigen-bait. For Spike immunizations, biotinylated RBD was employed as flow cytometric antigen-bait. Both these baits were biotinylated via their Avi-tag using a BirA500: BirA biotin-protein ligase standard reaction kit (Avidity). Baits were then pre-incubated for an hour on ice with streptavidin-APC (Biolegend 405207) or streptavidin-PE (Invitrogen 12-4317-87) prior to being added to the primary antibody stain.

### Flow cytometry

Popliteal LNs were dissected from mice and manually dissociated using Fisherbrand™ RNase-Free Disposable Pellet Pestles (Fisher Scientific 12-141-364) into FACS Buffer (2% FBS, 2 mM EDTA in 1x PBS). Cells were then incubated with Fc block (BD Biosciences 553142) and Live/Dead Zombie NIR™ (Biolegend 423105) in 1x PBS with 2 mM EDTA for 20 minutes on ice. Cells were subsequently stained with the primary antibody cocktail, including antigen-baits when necessary, for 20 minutes on ice. After staining, cells were washed 1x in FACS buffer and filtered using a Falcon™ Round-Bottom Polystyrene Test Tube with Cell Strainer Snap Cap (Falcon 08-771-23) prior to analysis on a BD FACSSymphony™ A3 or A5. For stains involving myeloid cells, lymph nodes were subjected to digestion with 200 U/mL of collagenase IV (STEMCELL 07427) and 20 ug/mL DNase I (STEMCELL 07900) in 1x HBSS in a 37°C water bath for 20 minutes and filtered via 100 µM strainers (Corning 431752) prior to analysis. All flow data was collected in BD FACSDiva™ and analyzed using FlowJo 10.10.0. For plots showing the percentage of PBs, population was calculated as a percentage of the Dump-channel.

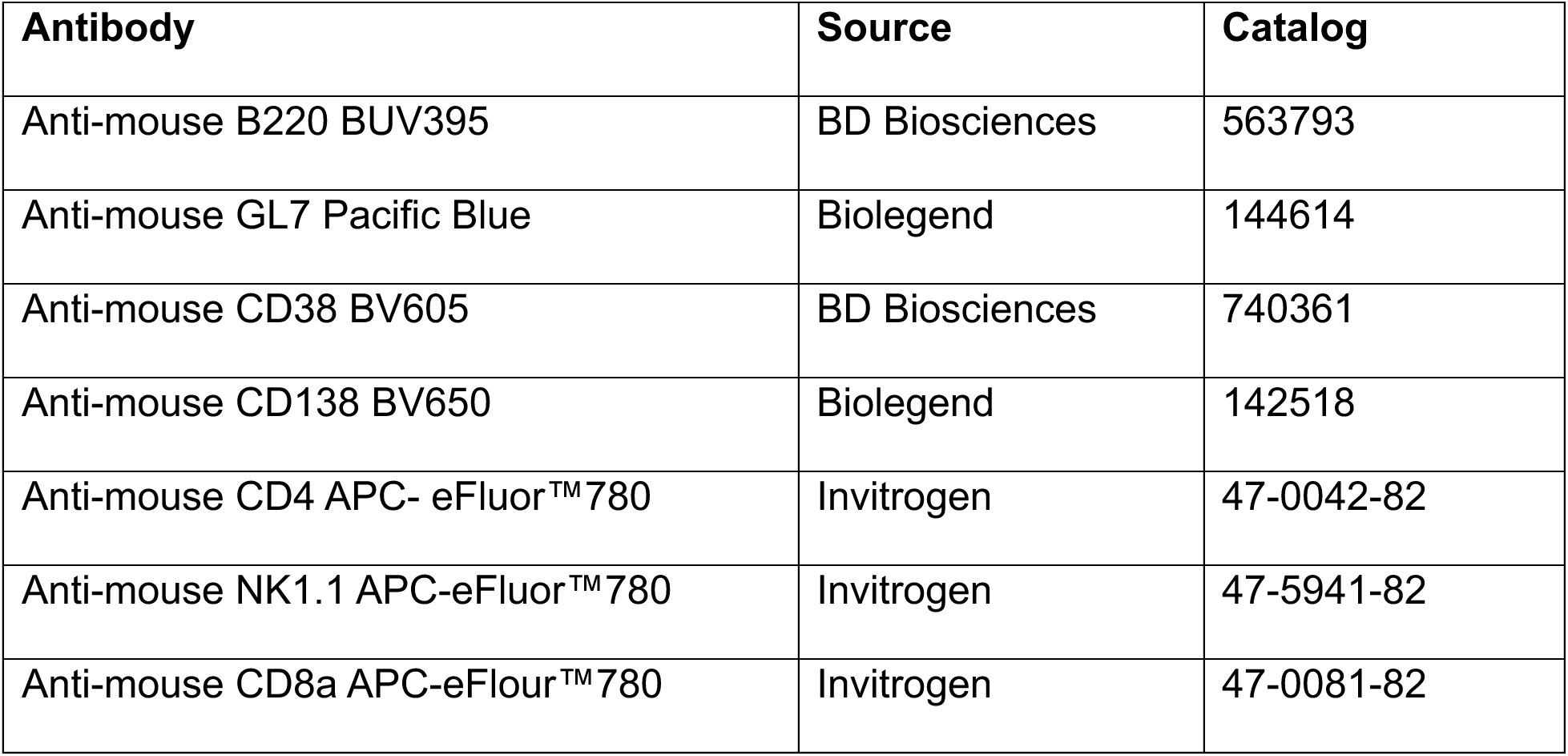
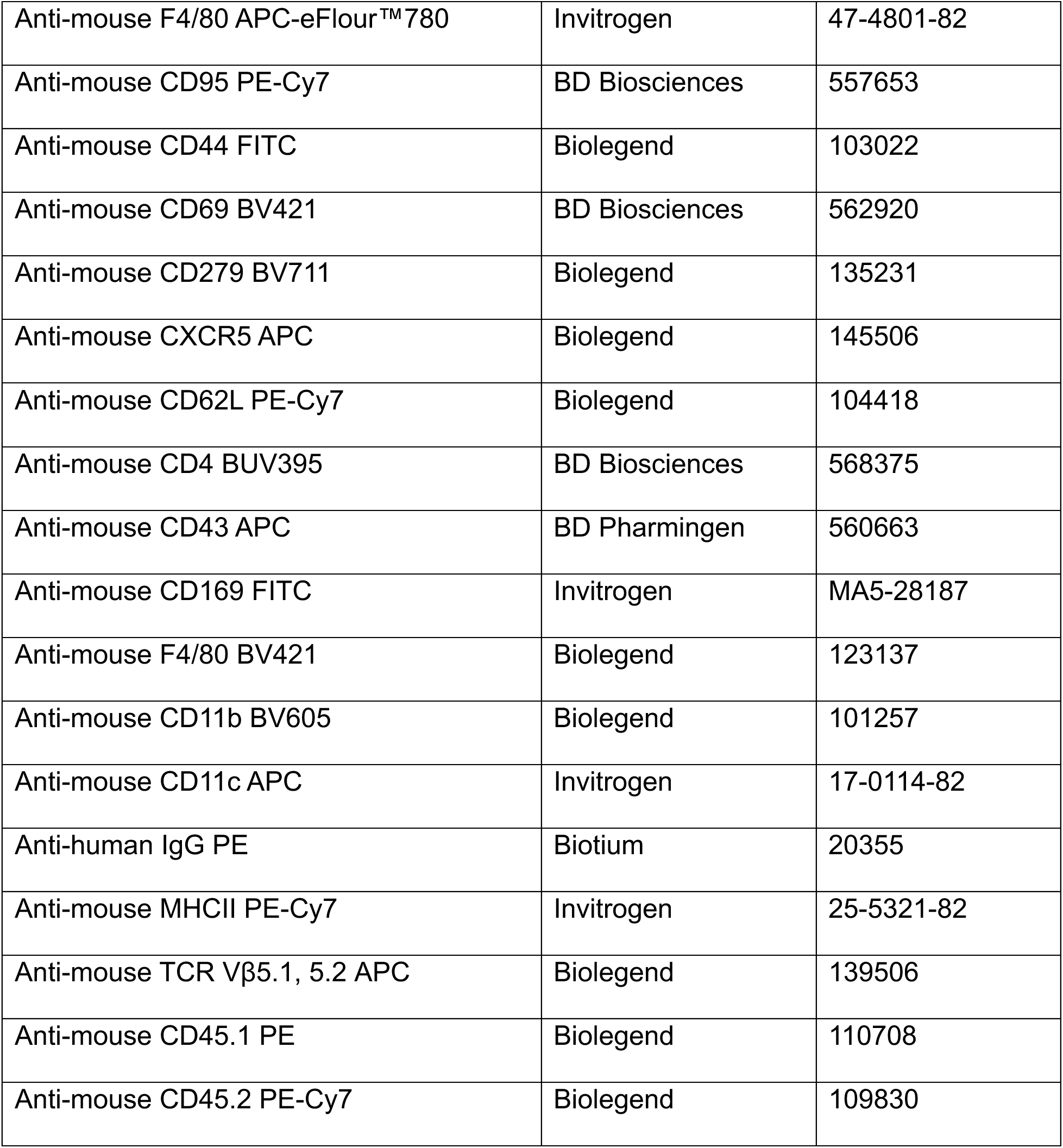

| Antibody | Source | Catalog |
| --- | --- | --- |
| Anti-mouse B220 BUV395 | BD Biosciences | 563793 |
| Anti-mouse GL7 Pacific Blue | Biolegend | 144614 |
| Anti-mouse CD38 BV605 | BD Biosciences | 740361 |
| Anti-mouse CD138 BV650 | Biolegend | 142518 |
| Anti-mouse CD4 APC- eFluor™780 | Invitrogen | 47-0042-82 |
| Anti-mouse NK1.1 APC-eFluor™780 | Invitrogen | 47-5941-82 |
| Anti-mouse CD8a APC-eFluor™780 | Invitrogen | 47-0081-82 |
| Anti-mouse F4/80 APC-eFlour™780 | Invitrogen | 47-4801-82 |
| Anti-mouse CD95 PE-Cy7 | BD Biosciences | 557653 |
| Anti-mouse CD44 FITC | Biolegend | 103022 |
| Anti-mouse CD69 BV421 | BD Biosciences | 562920 |
| Anti-mouse CD279 BV711 | Biolegend | 135231 |
| Anti-mouse CXCR5 APC | Biolegend | 145506 |
| Anti-mouse CD62L PE-Cy7 | Biolegend | 104418 |
| Anti-mouse CD4 BUV395 | BD Biosciences | 568375 |
| Anti-mouse CD43 APC | BD Pharmingen | 560663 |
| Anti-mouse CD169 FITC | Invitrogen | MA5-28187 |
| Anti-mouse F4/80 BV421 | Biolegend | 123137 |
| Anti-mouse CD11b BV605 | Biolegend | 101257 |
| Anti-mouse CD11c APC | Invitrogen | 17-0114-82 |
| Anti-human IgG PE | Biotium | 20355 |
| Anti-mouse MHCII PE-Cy7 | Invitrogen | 25-5321-82 |
| Anti-mouse TCR Vβ5.1, 5.2 APC | Biolegend | 139506 |
| Anti-mouse CD45.1 PE | Biolegend | 110708 |
| Anti-mouse CD45.2 PE-Cy7 | Biolegend | 109830 |

### Cell sorting and sequencing

For most cases, cells were processed as previously described (Viant et al., 2021). Briefly, single cells were sorted into 5 µL of 1% β-mercaptoethanol (Sigma-Aldrich M6250) in TCL buffer (Qiagen 1031576) in 96 well plates. RNA from these cells was then purified using RNAClean XP (Beckman Coulter A63987) and cDNA was generated using reverse Superscript III Reverse Transcriptase (Invitrogen 18080093). PCRs using previously defined primer combinations were then used to amplify heavy chain and kappa chain BCR sequences. PCR products underwent sanger sequencing and were then analyzed using the publicly available IgPipeline at the following URL: https://github.com/stratust/igpipeline/tree/igpipeline2_timepoint_v2. In the case of PBs sorted from FcRα null mice, cells were sorted into 384 well plates and underwent the SmartSeq3 protocol at the Sloan Kettering Institute Integrated Genomics Facility. SmartSeq3 libraries reads were trimmed using cutadapt v5.1 and aligned to the mouse GRCm38/mm10 genome build using STAR v2.7.11b with default parameters except for ‘--limitSjdbInsertNsj 2000000 --outFilterIntronMotifs –RemoveNoncanonicalUnannotated’ (Martin, 2011; Dobin et al., 2012). Reads were assigned with FeatureCounts, UMI reads were identified with the pattern ATTGCGCAATG, extracted and quantified with umi_tools (Liao et al., 2014; Hagemann-Jensen et al., 2020; Smith et al., 2017). BCR sequences were assembled using TRUST4 v1.1.7 (Song et al., 2021). Contigs containing less than 50 reads and more than one heavy or light chain were removed. Antibody heavy and light chains were paired and analyzed using igpipeline v.3.0 (https://github.com/stratust/igpipeline/tree/igpipeline3) with the mouse IMGT database as reference (Wang et al., 2023; Lefranc, 2011).

### Cell counts

Cell counts were either obtained using AccuCheck Counting Beads (Invitrogen PCB100) or via counting on a Cellometer K2 (Revvity CMT-K2-MX-150) with ViaStain™ AOPIStaining Solution (Revvity CS2-0106-5ML) and using FlowJo provided frequencies. Counts are rounded to the nearest hundredth.

### Cell transfers

For OT-II T cell transfer, cells were collected from spleens, mashed and filtered with a 100 µM filter (Corning 352360). Cells were subsequently treated with ACK (Gibco A1049201) to lyse red blood cells and T cells were isolated using the EasySep™ PE Positive Selection Kit II (STEMCELL 17684) with anti-CD4 PE (Biolegend 100512) according to manufacturer’s protocols. T cells obtained were then stained with CellTrace™ Violet Cell Proliferation Kit (Invitrogen C34557) for 20 minutes in a 37°C incubator and residual dye was quenched with 10% FBS in 1x PBS. Cells were washed twice in 1x PBS, counted on a Cellometer K2 and purity was confirmed via flow cytometry. 5 x 10^5^ T cells in 100-150 µL of PBS were injected intravenously into mouse recipients. For all OT-II experiments, transferred donor OT-II cells were congenically discordant at the CD45 locus compared to hosts (which were either WT CD45.1 B6 or WT C57BL6/J) allowing for identification of transferred cells.

### Bio-layer Interferometry

All BLI experiments were performed using a ForteBio Octet 96RedE at 1,000 RPM and 30°C. Fabs were diluted in 1x Kinetics Buffer (Satorius 18-1105) at a concentration of 35 µg/mL. Fabs were loaded onto FAB2G biosensors (Satorius 18-5125). TM4-Core was diluted to 100 µg/mL in 1x Kinetics Buffer. All samples were placed in 96-well plates (Greiner Bio-One 655209) in 200 µL volumes. Steps were as follows: (1) Baseline: 60s immersion of biosensors in 1x Kinetics Buffer, (2) Loading: 200s loading of Fabs onto biosensors, (3) Baseline2: 200s immersion of biosensors into 1x Kinetics Buffer, (4) Association: 300s immersion of biosensors with TM4-Core, (5) Dissociation: 600s immersion of biosensors back in 1x Kinetics Buffer. To normalize loading of the Fab, reference sensor data for each Fab was collected in the absence of TM4-Core and subtracted from traces obtained in the presence of TM4-Core. All analysis was performed using ForteBio software. Quality fit was validated both by visual checks and R^2^ values.

### Confocal microscopy

Microscopy was performed as previously described (MacLean et al., 2024). Briefly, popliteal LNs were collected, fixed in 4% PFA (Thermo Scientific 28908) for 2 hours, and subsequently dehydrated overnight in 30% sucrose. The following day LNs were washed in 1x PBS and embedded into Tissue-Tek® Cryomold® casts (Andwin No.NC9806558) with OCT (Fisher Healthcare 23-730-571). Lymph nodes were subsequently sliced on a cryostat to obtain ∼7 µM slices. For staining of slices, blocking was performed for 1 hour using Fc block (BD Biosciences 553142) and 5% normal mouse serum (Rockland B308) in 1x PBS containing .3% Triton X-100, .2% BSA and .1% sodium azide. Slides were then stained in 1x PBS containing .3% Triton X-100, .2% BSA, .1% sodium azide, 1% normal mouse serum and the primary antibody mix overnight. Antibodies used include anti-human IgG (Biotium 20355), anti-CD21/35 (Biolegend 123426), anti-CD169 (Invitrogen MA5-28187) and anti-CD138 (Biolegend 142504). The following day cells were washed 2x with 1x PBS and mounted using Fluoromount-G™ (Thermo Fisher 00-4958-02). Slides were visualized using a LSM 980 laser scanning confocal microscope (Zeiss). All analysis was performed using Imaris (Oxford Instruments, version 11.0). Due to their filamentous phenotype, the Surfaces function in Imaris was used to define FDC cell boundaries based on fluorescence intensity of CD21/35 staining. Statistics of mean fluorescence intensity of human 3BNC117 ICs using anti-human IgG on individual FDCs are reported. For quantification of ICs on non-CD21/35+ surfaces, the surfaces function in Imaris imaging software was similarly used. Surfaces were defined using the anti-human IgG channel, and surfaces that contained a mean intensity of CD21/35 above a minimum threshold were excluded. Statistics of mean fluorescence intensity of anti-human IgG on non-CD21/35+ surfaces are reported. All surfaces were manually verified.

### Statistical analysis

All statistical analysis performed is described in the figure legends. Analysis was performed using GraphPad Prism (version 10.2.3) and all figures were assembled using Adobe Illustrator.

## References

Bournazos, S., F. Klein, J. Pietzsch, M.S. Seaman, M.C. Nussenzweig, and J.V. Ravetch. 2014. Broadly Neutralizing Anti-HIV-1 Antibodies Require Fc Effector Functions for In Vivo Activity. Cell. 158:1243–1253. doi:10.1016/j.cell.2014.08.023.

Bournazos, S., and J.V. Ravetch. 2017. Anti-retroviral antibody FcγR-mediated effector functions. Immunol. Rev. 275:285–295. doi:10.1111/imr.12482.

Brüggemann, M., and K. Rajewsky. 1982. Regulation of the antibody response against hapten-coupled erythrocytes by monoclonal antihapten antibodies of various isotypes. Cell. Immunol. 71:365–373. doi:10.1016/0008-8749(82)90270-2.

Bruhns, P., B. Iannascoli, P. England, D.A. Mancardi, N. Fernandez, S. Jorieux, and M. Daëron. 2009. Specificity and affinity of human Fcγ receptors and their polymorphic variants for human IgG subclasses. Blood. 113:3716–3725. doi:10.1182/blood-2008-09-179754.

Dekkers, G., A.E.H. Bentlage, T.C. Stegmann, H.L. Howie, S. Lissenberg-Thunnissen, J. Zimring, T. Rispens, and G. Vidarsson. 2017. Affinity of human IgG subclasses to mouse Fc gamma receptors. mAbs. 9:767–773. doi:10.1080/19420862.2017.1323159.

Dobin, A., C.A. Davis, F. Schlesinger, J. Drenkow, C. Zaleski, S. Jha, P. Batut, M. Chaisson, and T.R. Gingeras. 2012. STAR: ultrafast universal RNA-seq aligner. Bioinformatics. 29:15–21. doi:10.1093/bioinformatics/bts635.

Dosenovic, P., E.E. Kara, A.-K. Pettersson, A.T. McGuire, M. Gray, H. Hartweger, E.S. Thientosapol, L. Stamatatos, and M.C. Nussenzweig. 2018. Anti–HIV-1 B cell responses are dependent on B cell precursor frequency and antigen-binding affinity. Proc. Natl. Acad. Sci. 115:4743–4748. doi:10.1073/pnas.1803457115.

Dvorscek, A.R., C.I. McKenzie, V.C. Stäheli, Z. Ding, J. White, S.A. Fabb, L. Lim, K. O’Donnell, C. Pitt, D. Christ, D.L. Hill, C.W. Pouton, D.L. Burnett, R. Brink, M.J. Robinson, D.M. Tarlinton, and I. Quast. 2024. Conversion of vaccines from low to high immunogenicity by antibodies with epitope complementarity. Immunity. 57:2433–2452.e7. doi:10.1016/j.immuni.2024.08.017.

Ellyard, J.I., D.T. Avery, T.G. Phan, N.J. Hare, P.D. Hodgkin, and S.G. Tangye. 2004. Antigen-selected, immunoglobulin-secreting cells persist in human spleen and bone marrow. Blood. 103:3805–3812. doi:10.1182/blood-2003-09-3109.

Enriquez-Rincon, F., and G.G. Klaus. 1984. Differing effects of monoclonal anti-hapten antibodies on humoral responses to soluble or particulate antigens. Immunology. 52:129–36.

Fang, Y., C. Xu, Y.-X. Fu, V.M. Holers, and H. Molina. 1998. Expression of Complement Receptors 1 and 2 on Follicular Dendritic Cells Is Necessary for the Generation of a Strong Antigen-Specific IgG Response. J. Immunol. 160:5273–5279. doi:10.4049/jimmunol.160.11.5273.

Finkelstein, M.S., and J.W. Uhr. 1964. Specific Inhibition of Antibody Formation by Passively Administered 19S and 7S Antibody. Science. 146:67–69. doi:10.1126/science.146.3640.67.

Getahun, A., J. Dahlström, S. Wernersson, and B. Heyman. 2004. IgG2a-Mediated Enhancement of Antibody and T Cell Responses and Its Relation to Inhibitory and Activating Fcγ Receptors. J. Immunol. 172:5269–5276. doi:10.4049/jimmunol.172.9.5269.

Gitlin, A.D., Z. Shulman, and M.C. Nussenzweig. 2014. Clonal selection in the germinal centre by regulated proliferation and hypermutation. Nature. 509:637–640. doi:10.1038/nature13300.

Hagemann-Jensen, M., C. Ziegenhain, P. Chen, D. Ramsköld, G.-J. Hendriks, A.J.M. Larsson, O.R. Faridani, and R. Sandberg. 2020. Single-cell RNA counting at allele and isoform resolution using Smart-seq3. Nat. Biotechnol. 38:708–714. doi:10.1038/s41587-020-0497-0.

Hägglöf, T., M. Cipolla, M. Loewe, S.T. Chen, L. Mesin, H. Hartweger, M.A. ElTanbouly, A. Cho, A. Gazumyan, V. Ramos, L. Stamatatos, T.Y. Oliveira, M.C. Nussenzweig, and C. Viant. 2023. Continuous germinal center invasion contributes to the diversity of the immune response. Cell. 186:147–161.e15. doi:10.1016/j.cell.2022.11.032.

Heesters, B.A., P. Chatterjee, Y.-A. Kim, S.F. Gonzalez, M.P. Kuligowski, T. Kirchhausen, and M.C. Carroll. 2013. Endocytosis and Recycling of Immune Complexes by Follicular Dendritic Cells Enhances B Cell Antigen Binding and Activation. Immunity. 38:1164–1175. doi:10.1016/j.immuni.2013.02.023.

Henry, C., and N.K. Jerne. 1968. Competition of 19S and 7S antigen receptors in the regulation of the primary immune response. J. Exp. Med. 128:133–152. doi:10.1084/jem.128.1.133.

Heyman, B. 1989. Inhibition of IgG-Mediated Immunosuppression by a Monoclonal Anti-Fc Receptor Antibody. Scand. J. Immunol. 29:121–126. doi:10.1111/j.1365-3083.1989.tb01106.x.

Inoue, T., and T. Kurosaki. 2024. Memory B cells. Nat. Rev. Immunol. 24:5–17. doi:10.1038/s41577-023-00897-3.

Kalergis, A.M., and J.V. Ravetch. 2002. Inducing Tumor Immunity through the Selective Engagement of Activating Fcγ Receptors on Dendritic Cells. J. Exp. Med. 195:1653– 1659. doi:10.1084/jem.20020338.

Kato, Y., R.K. Abbott, B.L. Freeman, S. Haupt, B. Groschel, M. Silva, S. Menis, D.J. Irvine, W.R. Schief, and S. Crotty. 2020. Multifaceted Effects of Antigen Valency on B Cell Response Composition and Differentiation In Vivo. Immunity. 53:548–563.e8. doi:10.1016/j.immuni.2020.08.001.

Klaus, G.G.B., M.B. Pepys, K. Kitajima, and B.A. Askonas. 1979. Activation of mouse complement by different classes of mouse antibody. Immunology. 38:687–695.

Künzli, M., and D. Masopust. 2023. CD4+ T cell memory. Nat. Immunol. 24:903–914. doi:10.1038/s41590-023-01510-4.

Lefranc, M.-P. 2011. IMGT, the International ImMunoGeneTics Information System. Cold Spring Harb. Protoc. 2011:pdb.top115. doi:10.1101/pdb.top115.

Liao, Y., G.K. Smyth, and W. Shi. 2014. featureCounts: an efficient general purpose program for assigning sequence reads to genomic features. Bioinformatics. 30:923– 930. doi:10.1093/bioinformatics/btt656.

MacLean, A.J., L.P. Deimel, P. Zhou, M.A. ElTanbouly, J. Merkenschlager, V. Ramos, G.S. Santos, T. Hägglöf, C.T. Mayer, B. Hernandez, A. Gazumyan, and M.C. Nussenzweig. 2024. Affinity maturation of antibody responses is mediated by differential plasma cell proliferation. Science. 387:413–420. doi:10.1126/science.adr6896.

Martin, M. 2011. Cutadapt removes adapter sequences from high-throughput sequencing reads. EMBnetJ. 17:10–12. doi:10.14806/ej.17.1.200.

Martínez-Riaño, A., S. Wang, S. Boeing, S. Minoughan, A. Casal, K.M. Spillane, B. Ludewig, and P. Tolar. 2023. Long-term retention of antigens in germinal centers is controlled by the spatial organization of the follicular dendritic cell network. Nat. Immunol. 1–14. doi:10.1038/s41590-023-01559-1.

McGuire, A.T., A.M. Dreyer, S. Carbonetti, A. Lippy, J. Glenn, J.F. Scheid, H. Mouquet, and L. Stamatatos. 2014. Antigen modification regulates competition of broad and narrow neutralizing HIV antibodies. Science. 346:1380–1383. doi:10.1126/science.1259206.

McNamara, H.A., A.H. Idris, H.J. Sutton, R. Vistein, B.J. Flynn, Y. Cai, K. Wiehe, K.E. Lyke, D. Chatterjee, N. KC, S. Chakravarty, B.K.L. Sim, S.L. Hoffman, M. Bonsignori, R.A. Seder, and I.A. Cockburn. 2020. Antibody Feedback Limits the Expansion of B Cell Responses to Malaria Vaccination but Drives Diversification of the Humoral Response. Cell Host Microbe. 28:572–585.e7. doi:10.1016/j.chom.2020.07.001.

Molina, H., V.M. Holers, B. Li, Y. Fung, S. Mariathasan, J. Goellner, J. Strauss-Schoenberger, R.W. Karr, and D.D. Chaplin. 1996. Markedly impaired humoral immune response in mice deficient in complement receptors 1 and 2. Proc. Natl. Acad. Sci. 93:3357–3361. doi:10.1073/pnas.93.8.3357.

Murgita, R.A., and S.I. Vas. 1972. Specific antibody-mediated effect on the immune response. Suppression and augmentation of the primary immune response in mice by different classes of antibodies. Immunology. 22:319–31.

Muta, T., T. Kurosaki, Z. Misulovin, M. Sanchez, M.C. Nussenzweig, and J.V. Ravetch. 1994. A 13-amino-acid motif in the cytoplasmic domain of FcγRIIB modulates B-cell receptor signalling. Nature. 368:70–73. doi:10.1038/368070a0.

Neuberger, M.S., and K. Rajewsky. 1981. Activation of mouse complement by monoclonal mouse antibodies. Eur. J. Immunol. 11:1012–1016. doi:10.1002/eji.1830111212.

Nimmerjahn, F., P. Bruhns, K. Horiuchi, and J.V. Ravetch. 2005. FcγRIV: A Novel FcR with Distinct IgG Subclass Specificity. Immunity. 23:41–51. doi:10.1016/j.immuni.2005.05.010.

Nimmerjahn, F., and J.V. Ravetch. 2005. Divergent Immunoglobulin G Subclass Activity Through Selective Fc Receptor Binding. Science. 310:1510–1512. doi:10.1126/science.1118948.

Pelletier, N., M. Casamayor-Pallejà, K.D. Luca, P. Mondière, F. Saltel, P. Jurdic, C. Bella, L. Genestier, and T. Defrance. 2006. The Endoplasmic Reticulum Is a Key Component of the Plasma Cell Death Pathway. J. Immunol. 176:1340–1347. doi:10.4049/jimmunol.176.3.1340.

Phan, T.G., I. Grigorova, T. Okada, and J.G. Cyster. 2007. Subcapsular encounter and complement-dependent transport of immune complexes by lymph node B cells. Nat. Immunol. 8:992–1000. doi:10.1038/ni1494.

Pincetic, A., S. Bournazos, D.J. DiLillo, J. Maamary, T.T. Wang, R. Dahan, B.-M. Fiebiger, and J.V. Ravetch. 2014. Type I and type II Fc receptors regulate innate and adaptive immunity. Nat. Immunol. 15:707–716. doi:10.1038/ni.2939.

Pincus, C.S., and V. Nussenzweig. 1969. Passive Antibody may simultaneously suppress and stimulate Antibody Formation against Different Portions of a Protein Molecule. Nature. 222:594–596. doi:10.1038/222594a0.

Poel, C.E. van der, G. Bajic, C.W. Macaulay, T. van den Broek, C.D. Ellson, G. Bouma, G.D. Victora, S.E. Degn, and M.C. Carroll. 2019. Follicular Dendritic Cells Modulate Germinal Center B Cell Diversity through FcγRIIB. Cell Rep. 29:2745–2755.e4. doi:10.1016/j.celrep.2019.10.086.

Qin, D., J. Wu, K.A. Vora, J.V. Ravetch, A.K. Szakal, T. Manser, and J.G. Tew. 2000. Fcγ Receptor IIB on Follicular Dendritic Cells Regulates the B Cell Recall Response. The J. Immunol. 164:6268–6275. doi:10.4049/jimmunol.164.12.6268.

Radoux, D., C. Kinet-Denoël, E. Heinen, M. Moeremans, J.D. Mey, and L.J. Simar. 1985. Retention of Immune Complexes by Fc Receptors on Mouse Follicular Dendritic Cells. Scand. J. Immunol. 21:345–353. doi:10.1111/j.1365-3083.1985.tb01440.x.

Ripperger, T.J., and D. Bhattacharya. 2021. Transcriptional and Metabolic Control of Memory B Cells and Plasma Cells. Annu Rev Immunol. 39:1–24. doi:10.1146/annurev-immunol-093019-125603.

Schaefer-Babajew, D., Z. Wang, F. Muecksch, A. Cho, M. Loewe, M. Cipolla, R. Raspe, B. Johnson, M. Canis, J. DaSilva, V. Ramos, M. Turroja, K.G. Millard, F. Schmidt, L. Witte, J. Dizon, I. Shimeliovich, K.-H. Yao, T.Y. Oliveira, A. Gazumyan, C. Gaebler, P.D. Bieniasz, T. Hatziioannou, M. Caskey, and M.C. Nussenzweig. 2023. Antibody feedback regulates immune memory after SARS-CoV-2 mRNA vaccination. Nature. 613:735–742. doi:10.1038/s41586-022-05609-w.

Scheid, J.F., H. Mouquet, B. Ueberheide, R. Diskin, F. Klein, T.Y.K. Oliveira, J. Pietzsch, D. Fenyo, A. Abadir, K. Velinzon, A. Hurley, S. Myung, F. Boulad, P. Poignard, D.R. Burton, F. Pereyra, D.D. Ho, B.D. Walker, M.S. Seaman, P.J. Bjorkman, B.T. Chait, and M.C. Nussenzweig. 2011. Sequence and Structural Convergence of Broad and Potent HIV Antibodies That Mimic CD4 Binding. Science. 333:1633–1637. doi:10.1126/science.1207227.

Shen, F.W., Y. Saga, G. Litman, G. Freeman, J.S. Tung, H. Cantor, and E.A. Boyse. 1985. Cloning of Ly-5 cDNA. Proc. Natl. Acad. Sci. 82:7360–7363. doi:10.1073/pnas.82.21.7360.

Sinclair, N.R. 1969. Regulation of the Immune Response: I. Reduction in Ability of Specific Antibody to Inhibit Long-Lasting IgG Immunological Priming After Removal of the Fc Fragment. J Exp Med. 129:1183–201.

Sinclair, N.R.STC., R.K. Lees, and E.V. Elliott. 1968. Role of the Fc Fragment in the Regulation of the Primary Immune Response. Nature. 220:1048–1049. doi:10.1038/2201048a0.

Smith, P., D.J. DiLillo, S. Bournazos, F. Li, and J.V. Ravetch. 2012. Mouse model recapitulating human Fcγ receptor structural and functional diversity. Proc. Natl. Acad. Sci. 109:6181–6186. doi:10.1073/pnas.1203954109.

Smith, T. 1909. Active immunity produced by so-called balanced or neutral mixtures of diphtheria toxin and antitoxin. J. Exp. Med. 11:241–256. doi:10.1084/jem.11.2.241.

Smith, T., A. Heger, and I. Sudbery. 2017. UMI-tools: modeling sequencing errors in Unique Molecular Identifiers to improve quantification accuracy. Genome Res. 27:491–499. doi:10.1101/gr.209601.116.

Song, L., D. Cohen, Z. Ouyang, Y. Cao, X. Hu, and X.S. Liu. 2021. TRUST4: immune repertoire reconstruction from bulk and single-cell RNA-seq data. Nat. Methods. 18:627–630. doi:10.1038/s41592-021-01142-2.

Ståhl, T.Dí.D., and B. Heyman. 2001. IgG2a-Mediated Enhancement of Antibody Responses is dependent on FcRγ+ Bone Marrow-Derived Cells. Scand. J. Immunol. 54:495–500. doi:10.1046/j.1365-3083.2001.01000.x.

Tam, H.H., M.B. Melo, M. Kang, J.M. Pelet, V.M. Ruda, M.H. Foley, J.K. Hu, S. Kumari, J. Crampton, A.D. Baldeon, R.W. Sanders, J.P. Moore, S. Crotty, R. Langer, D.G. Anderson, A.K. Chakraborty, and D.J. Irvine. 2016. Sustained antigen availability during germinal center initiation enhances antibody responses to vaccination. Proc. Natl. Acad. Sci. 113:E6639–E6648. doi:10.1073/pnas.1606050113.

Tang, A.F., G. Enyindah-Asonye, and C.E. Hioe. 2021. Immune Complex Vaccine Strategies to Combat HIV-1 and Other Infectious Diseases. Vaccines. 9:112. doi:10.3390/vaccines9020112.

Tao, T.-W., and J.W. Uhr. 1966. Capacity of Pepsin-digested Antibody to inhibit Antibody Formation. Nature. 212:208–209. doi:10.1038/212208a0.

Tas, J.M.J., J.-H. Koo, Y.-C. Lin, Z. Xie, J.M. Steichen, A.M. Jackson, B.M. Hauser, X. Wang, C.A. Cottrell, J.L. Torres, J.E. Warner, K.H. Kirsch, S.R. Weldon, B. Groschel, B. Nogal, G. Ozorowski, S. Bangaru, N. Phelps, Y. Adachi, S. Eskandarzadeh, M. Kubitz, D.R. Burton, D. Lingwood, A.G. Schmidt, U. Nair, A.B. Ward, W.R. Schief, and F.D. Batista. 2022. Antibodies from primary humoral responses modulate the recruitment of naive B cells during secondary responses. Immunity. 55:1856–1871.e6. doi:10.1016/j.immuni.2022.07.020.

Viant, C., A. Escolano, S.T. Chen, and M.C. Nussenzweig. 2021. Sequencing, cloning, and antigen binding analysis of monoclonal antibodies isolated from single mouse B cells. Star Protoc. 2:100389. doi:10.1016/j.xpro.2021.100389.

Walker, J.G., and G.W. Siskind. 1968. Studies on the control of antibody synthesis. Effect of antibody affinity upon its ability to suppress antibody formation. Immunology. 14:21–8.

Wang, X., B. Cho, K. Suzuki, Y. Xu, J.A. Green, J. An, and J.G. Cyster. 2011. Follicular dendritic cells help establish follicle identity and promote B cell retention in germinal centers. J. Exp. Med. 208:2497–2510. doi:10.1084/jem.20111449.

Wang, Z., F. Muecksch, R. Raspe, F. Johannsen, M. Turroja, M. Canis, M.A. ElTanbouly, G.S.S. Santos, B. Johnson, V.A. Baharani, R. Patejak, K.-H. Yao, B.J. Chirco, K.G. Millard, I. Shimeliovich, A. Gazumyan, T.Y. Oliveira, P.D. Bieniasz, T. Hatziioannou, M. Caskey, and M.C. Nussenzweig. 2023. Memory B cell development elicited by mRNA booster vaccinations in the elderly. J. Exp. Med. 220:e20230668. doi:10.1084/jem.20230668.

Wardemann, H., S. Yurasov, A. Schaefer, J.W. Young, E. Meffre, and M.C. Nussenzweig. 2003. Predominant Autoantibody Production by Early Human B Cell Precursors. Science. 301:1374–1377. doi:10.1126/science.1086907.

Weisel, F., and M. Shlomchik. 2015. Memory B Cells of Mice and Humans. Annu Rev Immunol. 35:1–30. doi:10.1146/annurev-immunol-041015-055531.

Weng, N., Y. Araki, and K. Subedi. 2012. The molecular basis of the memory T cell response: differential gene expression and its epigenetic regulation. Nat. Rev. Immunol. 12:306–315. doi:10.1038/nri3173.

Wernersson, S., M.C.I. Karlsson, J. Dahlström, R. Mattsson, J.S. Verbeek, and B. Heyman. 1999. IgG-Mediated Enhancement of Antibody Responses Is Low in Fc Receptor γ Chain-Deficient Mice and Increased in FcγRII-Deficient Mice. J. Immunol. 163:618–622. doi:10.4049/jimmunol.163.2.618.

Wiersma, E.J. 1992. Enhancement of the Antibody Response to Protein Antigens by Specific IgG under Different Experimental Conditions. Scand. J. Immunol. 36:193–200. doi:10.1111/j.1365-3083.1992.tb03091.x.

Wiersma, E.J., P.G. Coulie, and B. Heyman. 1989. Dual Immunoregulatory Effects of Monoclonal IgG-Antibodies: Suppression and Enhancement of the Antibody Response. Scand. J. Immunol. 29:439–448. doi:10.1111/j.1365-3083.1989.tb01143.x.

Wiersma, E.J., M. Nose, and B. Heyman. 1990. Evidence of IgG-mediated enhancement of the antibody response in vivo without complement activation via the classical pathway. Eur. J. Immunol. 20:2585–2589. doi:10.1002/eji.1830201209.

Yoshida, K., T.K. van den Berg, and C.D. Dijkstra. 1993. Two functionally different follicular dendritic cells in secondary lymphoid follicles of mouse spleen, as revealed by CR1/2 and FcR gamma II-mediated immune-complex trapping. Immunology. 80:34–9.

